# Ampk alpha2 T172 Activation Dictates Exercise Performance and Energy Transduction in Skeletal Muscle

**DOI:** 10.1101/2025.09.22.677805

**Authors:** Ryan N. Montalvo, Xiaolu Li, Gina Many, Tyler J. Sagendorf, Qing Yu, Wenqing Shen, Nishikant Wase, A. Robert Burgardt, Marina A. Gritsenko, Matthew J. Gaffrey, A. Hemangi Bhonsle, Yuntian Guan, Xuansong Mao, Mei Zhang, Wei-Jun Qian, Zhen Yan

**Author notes:** Corresponding Author: Zhen Yan. Lead Contact: Ryan N. Montalvo.

## Abstract

AMPK (5′-AMP-activated protein kinase) is an energetic sensor for metabolic regulation and integration. Here, we employed CRISPR/Cas9 to generate non-activatable Ampkα knock-in (KI) mice with mutation of threonine 172 phosphorylation site to alanine, circumventing the limitations of previous genetic interventions that disrupt the protein stoichiometry. KI mice of Ampkα2, but not Ampkα1, demonstrated phenotypic changes with increased fat-to-lean mass, impaired endurance exercise capacity, and diminished mitochondrial maximal respiration and conductance in skeletal muscle. Integrated temporal multi-omic analysis (proteomics/phosphoproteomics/metabolomics) in skeletal muscle at rest and during exercise establishes a pleiotropic yet imperative role of Ampkα2 T172 activation for glycolytic and oxidative metabolism, mitochondrial respiration, and contractile function. Importantly, there is a significant overlap of skeletal muscle proteomic changes in Ampkα2 T172A KI mice with that of type 2 diabetic patients. Our findings suggest that Ampkα2 T172 activation is critical for exercise performance and energy transduction in skeletal muscle and may serve as a therapeutic target for type 2 diabetes.

## INTRODUCTION

AMP-activated protein kinase-5’ (AMPK) is a critical energy-sensing molecule within skeletal muscle that has several established roles in regulating glucose uptake, fatty acid oxidation, and overall metabolic plasticity at rest, during exercise, and in response to exercise training^1–5^. AMPK exists as a heterotrimeric protein complex composed of a catalytic α subunit (α1 or α2 isoforms), a scaffolding β subunit (β1 or β2 isoforms) and a regulatory γ subunit (γ1, γ2, or γ3 isoforms) and is stimulated by AMP and/or ADP binding to the γ subunit, resulting in a conformational change that exposes the catalytic α threonine 172 (T172) site for phosphorylation by upstream kinases (e.g., CaMKKβ, LKB1)^6–9^, leading to full activation. In this regard, AMPK functions as a bellwether of cellular energy status in skeletal muscle, providing a mechanism by which the bioenergetic demand of exercise can trigger an immediate response to enhance metabolic flux during exercise and induce adaptive modifications that improve functional capacity for future endeavors.

Genetic manipulations of Ampk through global or muscle specific deletion of the Ampk α1/α2, β1/β2, or γ3 gene or forced expression of kinase-dead mutant Ampk in mice have demonstrated Ampk’s function in the regulation of glucose uptake, oxidative metabolism, mitochondrial biogenesis and respiration, and exercise performance^10–20^. Further, Ampk has also been shown to regulate mitophagy and mitochondrial quality control in skeletal muscle in response to exercise training^21,22^. Interestingly, several reports with genetic ablation of the Ampk genes demonstrated that Ampkα2 has a greater role than Ampkα1 in regulating skeletal muscle metabolic function, given that Ampkα2 is the dominant isoform in skeletal muscle^23–25^. Several of these studies advocate for the necessity of Ampkα2 signaling for the metabolic regulation in skeletal muscle^26–29^, others contend that Ampkα2 is dispensable for enhanced fatty acid oxidation induced by exercise or muscle contraction^30,31^. These contradicting findings warrant further clarification on these key determinants of skeletal muscle function.

Genetic deletion of a gene may not only disrupt protein stoichiometry but also cause ill-defined off-target effects. These unintended changes may cause compensatory adaptations, leading to spurious interpretation of the outcome measures^32,33^. Indeed, previous combinations of deletion of the Ampkα1/2, β1/2, or γ1/2/3 genes all caused disruption protein stoichiometry in Ampk holoenzyme and its protein complexes^17–19,34^. Therefore, despite the valuable insights provided by these genetic models, many of the phenotypic changes observed are discreet, and the interpretations may be confounded^35^.

Here, we took advantage of CRISPR/Cas9-mediated gene editing to substitute the threonine 172 phosphorylation site for alanine (T172A) in the α1 and α2 subunits separately, creating two global non-activatable knock-in (KI) models, circumventing the limitations of previous genetic models by maintaining protein stoichiometry. This approach made it possible to definitively ascertain the role of Ampk activation via T172 phosphorylation in regulation of muscle contractile and metabolic function and exercise-induced adaptations. We performed comprehensive biochemical, metabolic, and physiological assessments to dissect the functional role(s) of Ampkα1 and Ampkα2 activation on mitochondrial bioenergetics, metabolic control, and exercise performance. Further, we employed an integrated multi-omic (proteomic, phosphoproteomic, and metabolomic) approach to identify the impacts of ablated Ampkα2 activation on functional and metabolic phenotypes in skeletal muscle at rest and during exercise. Our findings provide robust evidence that phosphorylation of Ampkα2 at T172 is imperative to the maintenance of metabolic energy transfer and bioenergetic machinery at baseline and the regulation of these pathways during exercise in skeletal muscle. These results lay a detailed road map for directing research toward pharmacological and behavioral therapies that target Ampk for treating chronic metabolic disease, such as type 2 diabetes.

## RESULTS

### Ampkα2 T172A KI mice manifest an altered metabolic phenotype and exercise capacity

We first validated successful CRISPR Cas/9 Ampkα T172A substitution (knock-in; KI) by DNA sequencing and western blot (**Figure 1a-d; Figure S2a/b**). In addition to reduced Ampk phosphorylation, skeletal muscles from Ampkα2 KI mice showed decreased phosphorylation of classical Ampk targets^36,37^ (e.g. p-Acc; p-Ulk1), further confirming the validity of the KI model (**Figure S1a-c).** We then sought to extensively characterize the metabolic phenotypes of these KI mice. Firstly, body composition analysis by echoMRI revealed slightly reduced lean mass and increased fat mass in Ampkα2 KI mice (**Figure 1e- g**). Subsequently, we performed glucose tolerance and insulin tolerance tests. The Ampkα2 KI mice showed slightly improved glucose clearance (AUC) and fasting glucose with no change in insulin tolerance (**Figure 1h; Figure S1d-g**). Further, plantaris muscle weight and heart weight were lower in α2 KI mice without differences in gastrocnemius or soleus muscle weights (**Figure S1h-k**). Mice were then placed in metabolic cages to evaluate whole-body metabolism over 24 hours. Both WT and Ampkα2 KI mice showed an increased respiratory exchange ratio (RER) in the dark cycle with normal circadian patterns, indicating preserved metabolic activity. However, total and ambulatory activity were significantly lower in Ampkα2 KI mice in the dark cycle (**Fig 1i-l; Figure S1l/m**), suggesting underlying impairments in muscle function. No significant differences were observed for Ampkα1 KI mice related to any of these outcome measures, suggesting minimal functional roles of Ampkα1 activation in skeletal muscle (**Figure S2c-p**).

**Figure 1:**
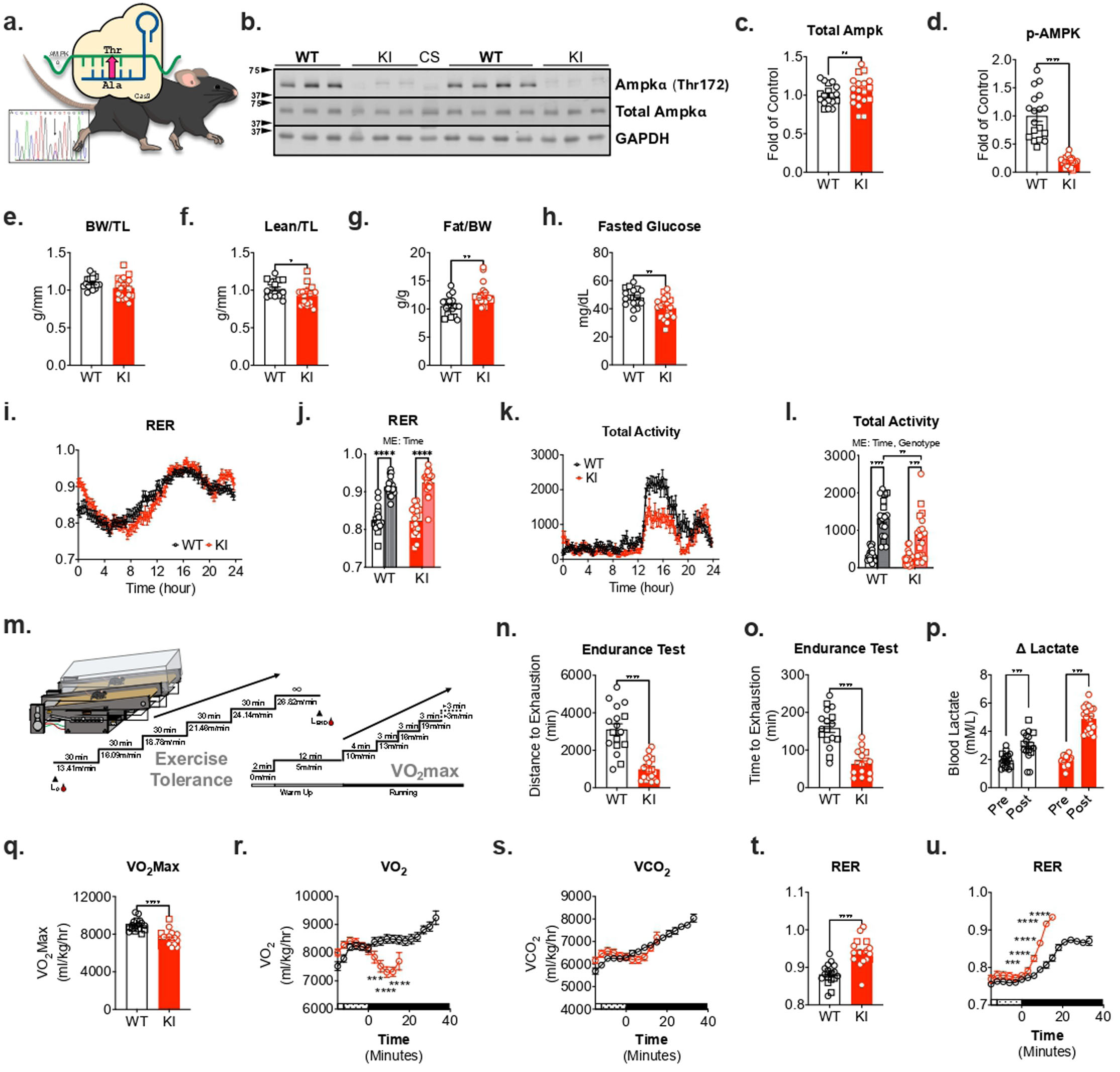
Ampkα2 KI mice demonstrate altered metabolic regulation and limited exercise capacity. Ampkα2 T712A KI and WT littermate mice were tested for genotype and phenotypic changes in body composition, metabolic function and exercise capacity. (**a-d**) Genotype was confirmed by DNA sequencing and western blot. WT= wild type, KI= Ampkα2 T172A KI, CS= common standard. (**f-g**) Body composition analysis (EchoMRI) for lean mass (g) normalized to tibia length (TL; mm) and fat mass (g) normalized to body weight (BW; g). (**h**) Blood glucose measurement was taken after an overnight fast (1700-0900). (**i-l**) Metabolic cage measurements (Columbus Instruments) were taken over 24 hours for respiratory exchange ratio (RER) and total activity. Comparisons made between the light/ resting (0700-1900; time 0-12 hours) and dark/ active (1900-0700; time 12-24 hours) cycles. (**m**) Illustration of protocols for endurance treadmill running and VO2Max tests for evaluation of exercise capacity. Blood collection for lactate before beginning (L0) and at the end (LEND) was indicated. (**n-p**) Endurance testing evaluated distance (meters) and time (minutes) and confirmed exhaustion through pre-post blood lactate measurements. (**q-u**) Gas exchange measurements were performed during VO2Max test for VO2Max, VCO2, and RER. Males (n=6 WT; n=7 KI) represented in squares and females (n=11 WT; n=12 KI) in circles for all outcomes; circles represent genotype average for **i/k/r/s/u**. Data presented as mean ± SEM. Statistical analysis performed by t-test between groups. Two-way ANOVA performed for **j/l** for main effects (ME) of time and genotype. Significance indicated as p < 0.05 (*), p < 0.01 (**), p < 0.001 (***), and p < 0.0001 (****), ns= not significant.

Various genetic deletions of Ampk genes in skeletal muscle demonstrate negative impacts on exercise capacity^11,12,16,27,38–40^, yet the disrupted protein stoichiometry may have convoluted these results. To this end, we measured exercise performance for Ampkα1 KI and Ampkα2 KI mice in treadmill running endurance test as well as a separate VO2max test to investigate the role of Ampk activation during exercise. The procedures for each exercise test are illustrated in **Figure 1m**. Ampkα2 KI mice had dramatically diminished time to fatigue during the endurance test compared to WT littermate mice (**Figure 1n-p**); this effect was not observed in Ampkα1 KI mice (**Figure S2q-s**). Further, VO2max was significantly diminished in Ampkα2 KI mice (**Figure q-u**). These results are further corroborated by impaired metabolic regulation represented by an early elevation of RER, which suggests aberrant fatty acid oxidation and a lower anaerobic threshold (**Figure 1t-u; Figure S1n-p).** Like the endurance test, no significant effects on VO2max outcomes were observed for the Ampkα1 KI mice (**Figure S 2t-v**). Taken together, these results demonstrate a deleterious effect of Ampkα2 T172 signaling ablation within the skeletal muscle, as is evident by significantly reduced functional capacity as well as impaired metabolic regulation.

### Global proteomics demonstrates prevalent regulatory role of Ampkα2 T172 activation in mitochondrial bioenergetic machinery

Our data demonstrate that Ampkα2 T172 activation dictates the gross functional capacity and metabolic signature in skeletal muscle at rest and during exercise, which is consistent with previous studies^41,42^; however, the molecular mechanisms underlying the of the role of Ampkα2 T172 activation has not been investigated. We thus conducted a global proteomic analysis of Ampkα2 KI skeletal muscle with two objectives: (1) to provide high-resolution mapping of the known roles of Ampkα2 in various regulatory processes, and (2) to identify novel targets of Ampkα2 activation that may impact exercise and skeletal muscle functional capacity.

Relative to WT mice, Ampkα2 KI mice displayed 444 significantly altered proteins in females and 1310 in males (adjusted p < 0.05). (**Supp. data file 1**). We next applied the pre-ranked version of the Correlation Adjusted MEan RAnk gene set test (CAMERA-PR)^43^ to identify concordant changes in Gene Ontology (GO) terms (Molecular Function (MF); Biological Processes (BP); Cellular Components (CC)) (**Figure S3a-c**). Ampkα2 KI mice had profound changes of proteins in mitochondrial and core metabolic pathways that include the electron transport chain, cellular respiration, TCA cycle, and ATP metabolic process (**Figure 2a-b**). Regarding metabolism-related proteins, male and female KI mice shared 76 differentially expressed proteins, 64 of which decreased in abundance (**Figure 2c**). Further, Ampkα2 KI mice showed significantly decreased electron transport and core metabolic proteins in skeletal muscle in both sexes (**Figure 2d**); although Ampkα2 KI in male mice had a greater impact on mitochondrial proteins compared to females. Importantly, pyruvate dehydrogenase (Pdh), which serves as an integration point between anaerobic and aerobic pathways, was decreased in both male and female Ampkα2 KI mice (**Fig 2d**). Male but not female α2 KI mice displayed reduced abundance of proteins related to pyruvate transport (Mpc1/Mpc2), and the Pdh subunits E1 (Pdha1, Pdhb, Pdhx, E2 (Dlat), and E3 (Dld)) as well as Pdh phosphatase (activating) (Pdpr; Pdp1) and Pdh kinase (inactivating) isozymes 2 and 4 (Pdk2; Pdk4). Pdh E3 simultaneously produces NADH and FADH2, the substrates of ETC complex I and II, respectively, demonstrating a dramatic effect of Ampkα2 T172 inactivation on Pdh activation in male mice. Male mice appear to be more susceptible to the disruption of this signaling pathway. These findings suggest that Ampkα2 activation in skeletal muscle plays a fundamentally important role in maintaining the metabolic machineries.

**Figure 2:**
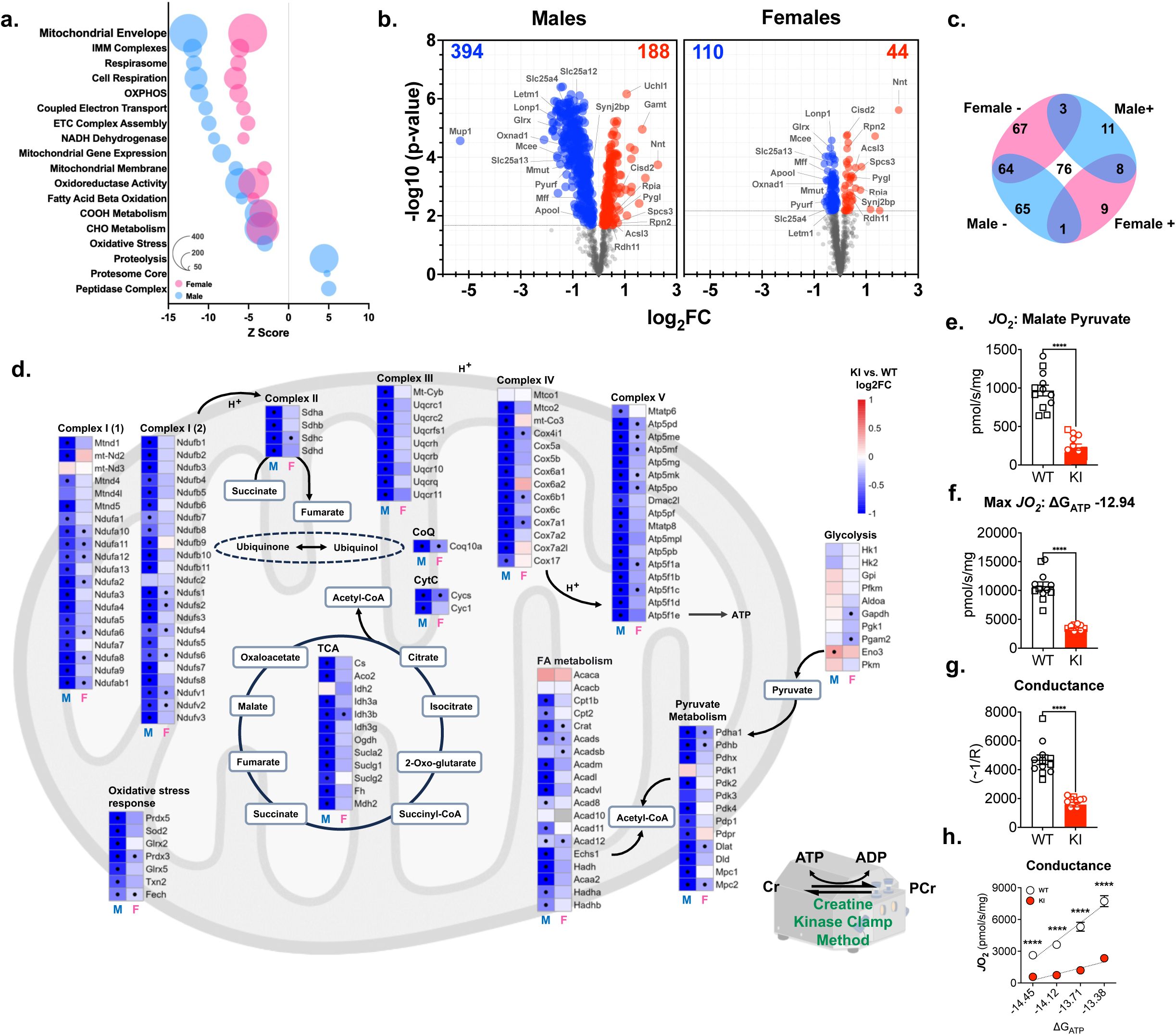
Global proteomic analysis reveals Ampkα2 KI mice have disrupted core metabolic machineries for mitochondrial function. Global proteomics was performed for gastrocnemius muscle from male and female WT (n=6) and Ampkα2 KI (n=6) mice. (**a**) Dot plot of the most significantly regulated GO terms (GO biological processes (BP); cellular components (CC) and molecular function (MF) represented by Z score (x-axis), number of proteins per GO term (size of dot), separated by males (blue) and females (pink) as indicated in legend. (**b**) Volcano plot of proteins represented in males (394 proteins significantly downregulated (blue) and 188 upregulated (red)) and females (110 downregulated and 44 upregulated) limited to include GO term gene sets designated in **a**. (**c**) 4-way representation of significantly regulated proteins that overlapped between males and female volcano plots; upregulated (+) and downregulated (-), 76 proteins in total (center). (**d**) enrichment analysis of proteins related to glycolysis and mitochondrial oxidative metabolism (pyruvate metabolism, TCA, fatty acid (FA) oxidation, oxidative stress response and electron transport chain complexes (I, II, III, IV, V); • representative of adjusted p < 0.05 difference within heatmap by t-test. (**e-h**) Creatine kinase clamp analysis of mitochondrial respiration with malate-pyruvate substrates in a non-phosphorylating substrate-stimulated state and maximal respiration (MAX *J*O2; ΔGATP -12.94). Conductance measured through PCr titrations of 1 mM (ΔGATP -13.38 kCal/mol), 2 mM (ΔGATP -13.71 kCal/mol), 4 mM (ΔGATP -14.12 kCal/mol), and 6 mM (ΔGATP -14.45 kCal/mol) as slope of ΔGATP and *J*O2. Males (n=6 WT; n=6 KI) are represented in squares and females in circles (n=6 WT; n=6 KI); average of genotypes represented in circles in **h**. Data presented as mean ± SEM. Statistical analysis performed by t-test between groups: p < 0.0001 (****).

To gain further insights into the regulatory role of Ampkα2 T172 in skeletal muscle mitochondrial processes, we performed gene set enrichment analysis utilizing the reference gene set MitoCarta3.0^43^ to identify mitochondria-specific alterations in protein abundance. This investigation revealed decrements of proteins related to mitochondrial genetic replication (i.e., TOM; TIM; RNA granule; ribosomes), mitochondrial and ETC assembly and integrity (i.e. MICOS; MIB complexes), and mitochondrial quality control (e.g., fusion, fission, mitophagy) (**Figure S4a-d**). Again, the decrements were more severe in male KI mice compared with female KI mice.

In an effort to directly dissect the role of Ampkα2 T172 activation on mitochondrial function, we measured mitochondrial respiration in isolated mitochondria from gastrocnemius. Further, we employ a creatine-kinase clamp protocol amended to include phosphocreatine titrations as a measure of mitochondrial conductance over a range of respiratory demands (ΔGATP) (detailed in **Figure S4e/f**)^44,45^. Substrate-stimulated non-phosphorylating respiration utilizing malate pyruvate and phosphorylating maximal respiration (MAX *J*O2; ΔGATP -12.94) were greatly decreased in Ampkα2 KI mice (**Figure 4e/f**). Measurement of conductance (slope of *J*O2 vs. ΔGATP -13.38, -13.71, -14.12, and -14.45) mimics a tractable bioenergetic stress test, and Ampkα2 KI mice demonstrated increased resistance within the electron transport chain and impaired ability to respond to energetic demand (**Figure 4g/h**). Although the roles of Ampk on mitochondrial quality and function have been implicated in several studies^16,17,21,22,46^, this is the first report to show direct evidence of the essential role of Ampkα2 T172 activation in maintaining normal mitochondrial machinery, respiration, and conductance in skeletal muscle.

### Exercise reveals a potential role of Ampkα2 T172 activation in skeletal muscle metabolic and contractile function via phosphoproteome regulation

Here we used phosphoproteomics to distinguish skeletal muscle signaling in response to treadmill running to identify novel targets of Ampk that may impact overall functional capacity and metabolic plasticity. Based on the early elevation of RER during VO2max testing, we collected samples at an intermediate 10-minute (10 min) timepoint and at exhaustion (EXH) to compare exercise responses to the sedentary states (protocol demonstrated in **Figure S5a; Supp. data file 2**). The phosphoproteomic response at 10 minutes was significantly greater in Ampkα2 KI mice compared with WT mice, which is consistent with the observations of reduced exercise capacity in Ampkα2 KI mice (**Figure S5b/c**). At exhaustion, male Ampkα2 KI mice had more changes compared to WT mice. Interestingly, female WT mice showed more changes than female Ampkα2 KI at exhaustion. Overall, female mice had significantly fewer changes compared to males, displaying a clear sex difference. To elucidate the functionally diverse phosphoproteomic response to acute exercise, we performed ingenuity pathway analysis (IPA), which revealed a variety of signaling mechanisms impacted by Ampkα2 T172 signaling ablation, including those related to insulin signaling, ERK/MAPK, mTOR, muscle structure and contraction, nNOS, apoptosis, and actin cytoskeleton (**Fig 3a; Supp. data file 3**).

**Figure 3:**
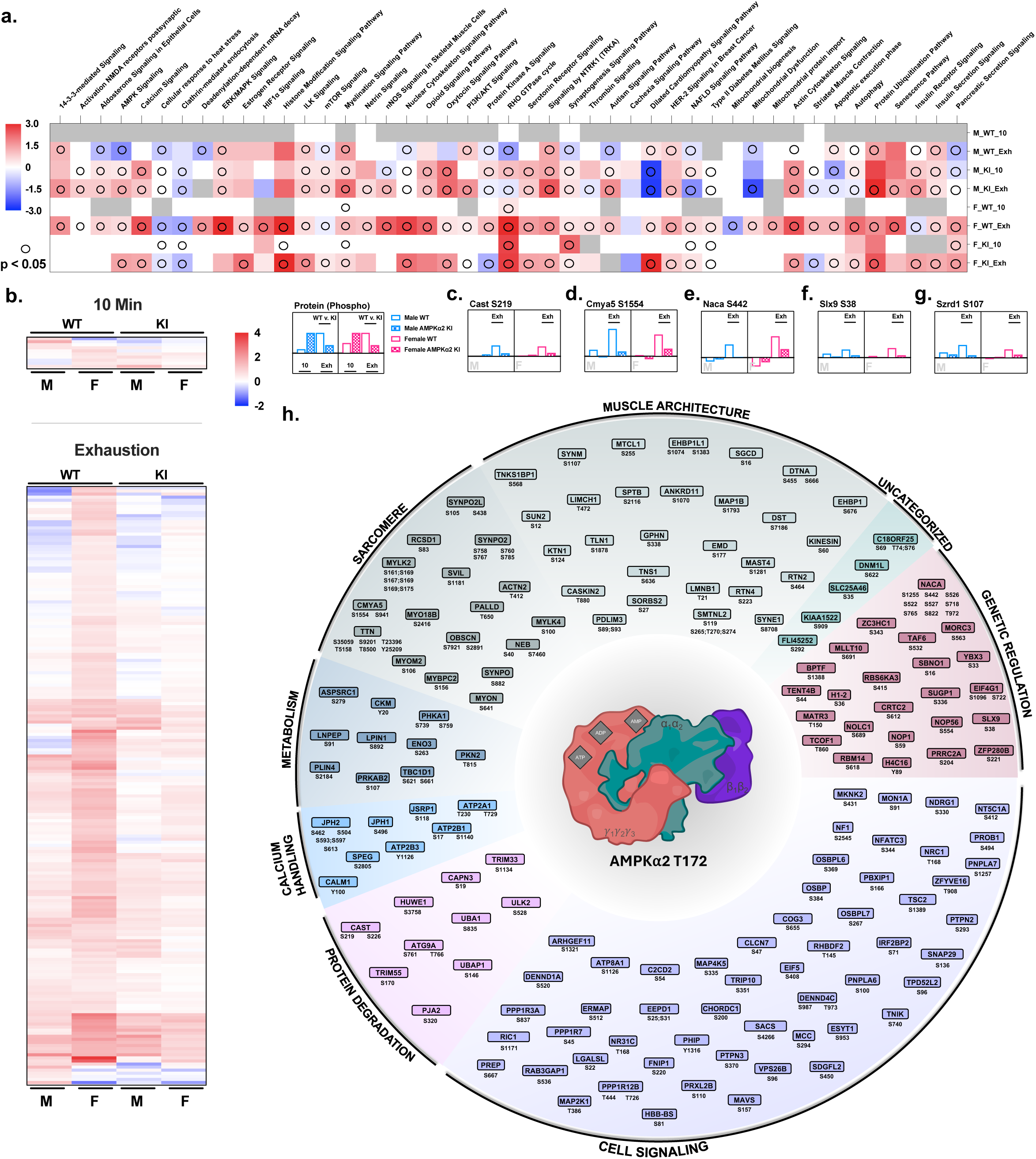
Ampkα2 T712 activation dictates the phosphoproteomic response to exercise in skeletal muscle. Male and female Ampkα2 KI and WT littermate mice were divided into three groups (1) sedentary, (2) exercise for 10 minutes (10 min) and (3) exercise to exhaustion (Exh) groups. Protocol detailed in figure S5. (**a**) Ingenuity Pathway Analysis (IPA) heat map of phosphorylation sites comparing male (M) and female (F) at 10 min (10) and exhaustion (Exh). Empty circle O indicates the significance of the pathway with p< 0.05. (**b**) putative phosphorylation sites significantly activated (adjusted p< 0.05) by exercise but not activated in male female KI mice at 10 min (top) and exhaustion (bottom). Data organized by hierarchical clustering: Euclidean complete. (**c-g**) Log2FC of phosphorylation sites representing significantly increased phosphorylation in male (blue bar) and female (pink bar) Ampkα2 KI (checkered bar) and WT (open bar) mice. A significant decrease of phosphorylation was observed in Ampkα2 KI (p<0.05) at exhaustion for Calpastatin (*Cast*) S219, cardiomyopathy associated protein 5 (*Cmya5*) S1554, nascent polypeptide associated complex subunit alpha (*Naca*) S442, SLX9 ribosome biogenesis factor (*Slx9*) S38, and SUZ RNA binding domain containing 1 (*Szrd1*) S107. Bars above timepoints indicate significant differences between WT and Ampkα2 KI mice at 10 minutes or exhaustion. (**h**) all potential sites identified as under regulation by Ampkα2 activation at 10 min or exhaustion designated into functional groups irrespective of sex.

The main goal of this analysis is to identify putative substrates of activated Ampkα2 via T172 phosphorylation by exercise. To this end, we compared fold change of phosphorylation at various phosphorylate sites over the sedentary condition for both genotypes separately at the 10 minute and at exhaustion with results limited to those with adjusted p < 0.05. Further, we compared WT with Ampkα2 KI mice to observe significant differences at these separate timepoints (i.e., Δ log2FC; **Supp. data file 2**) with consideration of sex. This analysis revealed 10 unique phosphorylation sites that were significantly regulated at the 10 min time point and 194 at exhaustion (**Figure 3b**). Interestingly, only 5 sites were significantly increased at exhaustion in WT mice that were also were significantly diminished in Ampkα2 KI for both the males and females: Calpastatin (Cast) S219, cardiomyopathy associated protein 5 (Cmya5, [Myospryn]) S1554, nascent polypeptide associated complex subunit alpha (Naca) S442, SLX9 ribosome biogenesis factor (Slx9) S38, SUZ RNA binding domain containing 1 (Szrd1) S107 (**Figure 3c- g**). We consider these novel targets of Ampkα2.

Further evaluation of the significantly regulated phosphorylation sites allowed for binning of genes into eight physiologically relevant groups (**Figure 3h**): muscle architecture, sarcomere, metabolism, genetic regulation, general cell signaling, protein degradation, calcium handling, and uncategorized cellular processes. Although the specific role for many of these phosphorylation sites have yet to be determined, these genes indicate a significant regulatory control of Ampkα2 activation via T172 phosphorylation on the functional process of excitation contraction coupling, muscle structural stability and contractile capacity and metabolism (**Supp.** Figure 6). Firstly, **calcium handling** at T-tubule ryanodine receptor (e.g., Junctophillin 1/2 (Jph1/ Jph2), sarcoplasmic reticulum calcium ATPase 1 (Atp2a1), and intracellularly (e.g., calmodulin (Calm1)) are significantly altered. Additional changes were noted in **sarcomere** and specifically at Z disk (e.g., Cmya5; synaptotodin (Synpo); supervilin (Svil); Titin (Titin)), which is complemented by alterations in exercise-responsive phosphorylation sites related to gross **muscle architecture** (e.g., sarcoglycan delta (Scgd) and dystonin (Dst)). Changes in sarcomeric protein phosphorylation are consistent with global proteomics results, suggesting increased phosphorylation of proteins related to **protein degradation** in response to exercise via the ubiquitin proteasome system (e.g., Cast; calpain-3 (Capn3)). Of note, our results substantiate recent reports that identified C18orf25^47^ and Fnip1^48^ as novel Ampk targets and essential regulators of exercise capacity.

The role of Ampkα2 activation in **metabolism** is well-demonstrated in the current analysis, primarily in glucose regulation. We observed significant phosphorylation of Ampk upstream regulator, serine/threonine-protein kinase N2 (Pkn2), which has been shown to considerably impact glucose metabolism through Ampk activation in skeletal muscle^49^. Glycolytic regulation and glucose handling are also highlighted with significant results in Tbc1d1 and enolase (Eno3) as well as glycogen metabolism via phosphorylase kinase alpha (Phka1). Further cytosolic muscle specific creatine kinase (Ckm Y20) was phosphorylated at the 10 minutes and exhaustive timepoints for male WT mice but not in the Ampkα2 T172A KI mice. Ckm is essential for the metabolic response to exercise and metabolic flexibility within skeletal muscle, providing a novel and substantial regulatory role of Ampkα2 T172 activation.

Further experiments will be required to elucidate Ampk’s integrative regulation of skeletal muscle architecture, contractile capacity and energetic supply. Nevertheless, the current study provides evidence that Ampkα2 T172 activation synchronizes skeletal muscle metabolic function and contractile capacity through coordinated regulation metabolic and contractile mechanisms during exercise.

### Ampkα2 T172 activation dictates glycolytic and oxidative flux during exercise

We then performed a temporal metabolomic analysis to help reconcile our phosphoproteomic and proteomic findings and ascertain metabolic regulation in Ampkα2 KI mice (**Figure 4a; Supp. data file 4**). At 10 minutes, an irregular response in the metabolites relating to glucose 1 phosphate (G1P; **Figure 4b**), trehalose), L-aspartic acid, and fatty acid metabolism (e.g., L-carnitine; **Figure 4c**) was observed in male Ampkα2 KI mice. Decreased levels of metabolites indicates a more rapid use and reliance on these pathways for energy production in the absence of Ampkα2 T172 activation. Contrastingly, at 10 minutes some metabolites increased in abundance in male KI mice which included adenine (**Figure 4d**), adenosine 3’,5’ diphosphate, and pyruvate (**Figure 4e**), indicating decreased utilization of these substrates. These metabolites and NAD^+^ (**Figure 4f**) were also increased at exhaustion in Ampkα2 KI mice, demonstrating a more wide-spread impairment of these metabolic pathways. Of note, 4-hydroxy- L-proline, L-kynurenine, and methylmalonic acid (**Figure 4g**) were significantly increased at exhaustion in Ampkα2 KI but not WT mice. Interestingly, WT mice but not Ampkα2 KI mice showed significantly reduced propionic acid level at exhaustion, highlighting the importance of Ampka2 activation in anaplerotic metabolism. Male KI mice further demonstrated a heightened oxidative environment with increased level of oxidized glutathione at exhaustion (**Figure 4h**), which may potentially impair metabolic flux and contribute to sarcomere instability.

**Figure 4:**
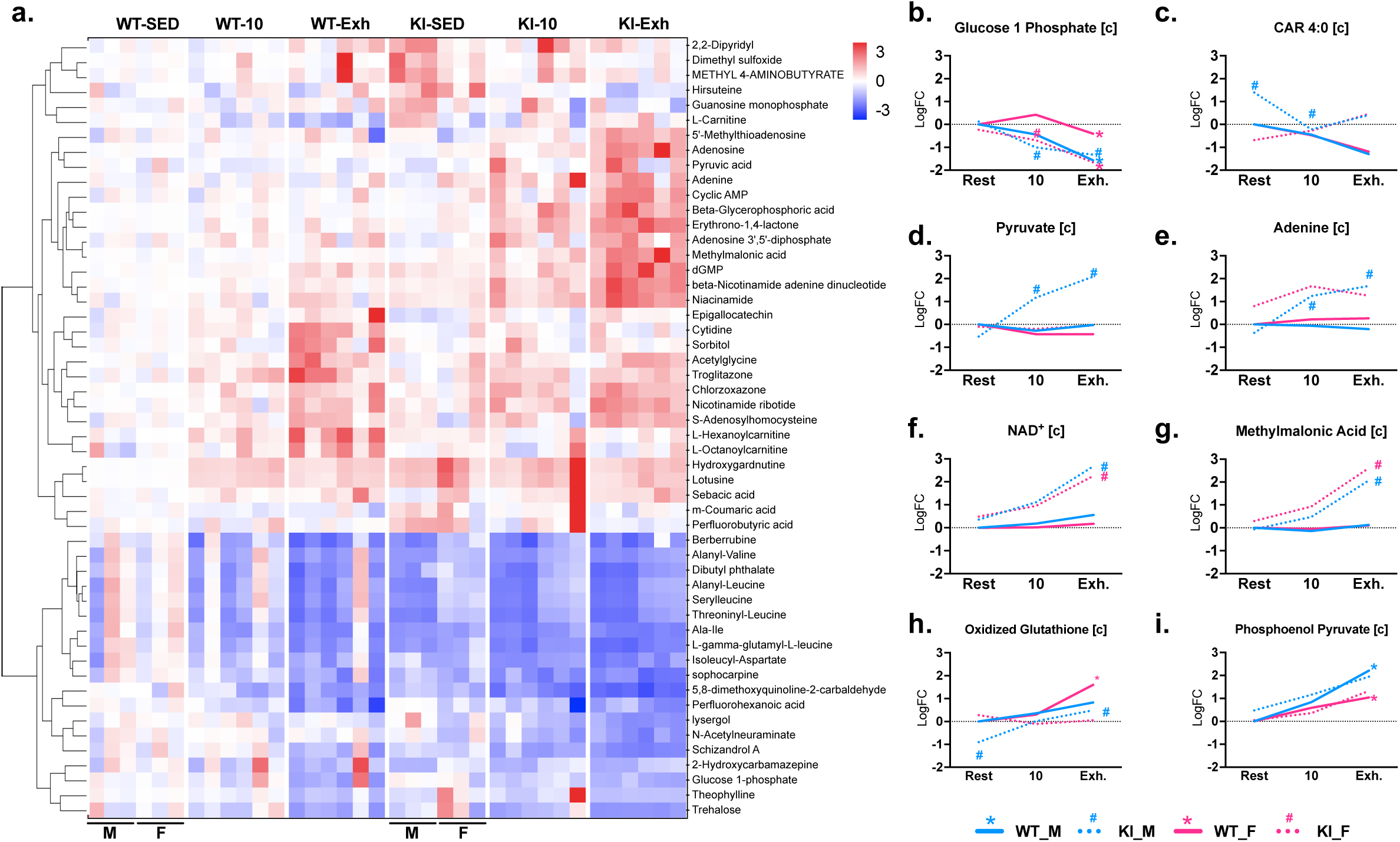
Metabolomic response to exercise demonstrates increased reliance on glycolysis in Ampkα2 KI mice. Metabolomic analysis was performed (protocol detailed Figure S5) for gastrocnemius muscle for Ampkα2 KI and WT mice (WT n=6; KI n=6) in sedentary state (SED), at 10 min of (10) exercise and at exhaustion (Exh). (**a**) heat map of data from male (M) and female (F) mice; organized by Euclidean complete hierarchical clustering (p < 0.01), individual metabolites listed to right of heatmap. (**b-i**) representative tracings of male (blue) and female (pink) WT (solid line) and Ampkα2 KI (dotted line) metabolites during the exhaustive exercise protocol. Adjusted p < 0.05 indicated by colored * (WT) or ^#^ (KI) at rest (0), 10 min (10), and exhaustion (Exh).

WT and Ampkα2 KI mice show decreased G1P at exhaustion while male KI mice show this change at 10 minutes of exercise. A similar pattern was also observed for carbohydrate trehalose. These findings suggest a more severe dysregulation of glycolytic metabolism in male Ampkα2 KI mice with exercise. Phosphoenolpyruvic acid (**Figure 4i**) was increased in both WT (adjusted p < 0.05) and Ampkα2 KI mice (adjusted p = 0.10 M, 0.09F) at exhaustion, along with increased pyruvate at 10 and 90 minutes of exercise for male KI mice. These findings are line with the observed increase of enolase 3 expression, suggesting increased glycoltyic flux for male Ampkα2 KI mice (**Figure 2d**). Lastly, WT mice demonstrate a decrease in lactic acid without a significant change in Ampkα2 KI mice. Agreement amongst sex is present in many of these results but the accumulation of pyruvate, adenine, and oxidized glutathione in male Ampkα2 KI mice at exhaustion suggest a sex-specific aberrant impact. Our integrative analysis highlights that Pdh kinase 2 and 4 (inactivating) are positively regulated by increased ATP:ADP and NADH:NAD^+^, which were diminished in WT but not Ampkα2 KI mice during exercise (**Figure 2d**; **Figure 4e/f**). These results suggest that Ampkα2 KI mice initiate glycolysis more rapidly and are more reliant on this processs during exercise, contrasted by a decreased flux through the TCA cycle. This notion is in line with our VO2max testing, mitochondrial respiratory outcomes, and phosphoproteomics analysis.

### Significant overlap of skeletal muscle proteomic features between Ampkα2 KI mice and human diabetic patients

Ampk could be an important target for mitigating obesogenic pathology in skeletal muscle, but not enough is understood about the role of Ampkα2 T172 activation in the context of type 2 diabetes (T2D)^41,42^. To this end, we investigated the overlap of the current data set with that of skeletal muscles of diabetic patients to identify specific targets of Ampkα2 T172 activation that may serve to mitigate diabetic pathology.

We first plotted the coordinated overlap of our global proteomics results with two proteomic data sets from skeletal muscles of diabetic patients^50,51^ (**Figure S6; Fig 5a**). Proteins that share decreased expression between diabetic patients and Ampkα2 KI mice (q3) may unveil the putative role of reduced Ampkα2 T172 in T2D. We observed a profound correlation between T2D patients and Ampkα2 KI mice that elucidates a reduced expression in the regulation of mitochondrial energy transfer: substrate utilization, transport, and electron transmission (e.g., Slc25a11 (malate aspartate shuttle) and electron transfer flavoproteins Etfa/ Etfb)), electron potential energy generation through NADH and FADH2 (e.g., Acyl-CoA dehydrogenase medium (Acadm); isocitrate dehyrogenase (Idh3a), ETC complexes (e.g., Ndufv1 (C1); Sdha (CII); Uqcr10 (CIII); Cox7a2 (CIV)), and maintenance of phosphorylation potential (e.g., adenine nucleotide translocase (Ant1); phosphate carrier (Slc25a3); mitochondrial creatine kinase (Ckmt2)) (**Figure 5a**). Further, we identified 17 proteins that were present in male and female Ampkα2 KI as well as both T2D studies: Acls1; Ckmt2; Cox4i1; Cycs; Dlat; Hsd17b10; Hsdl2; Hspe1; Ndufa10; Ndufa6; Ndufa8; Ndufab1; Ndufs6; Ndufv1; Nnt; Slc25a12; Slc25a4 (annotated in purple and listed in table).

**Figure 5:**
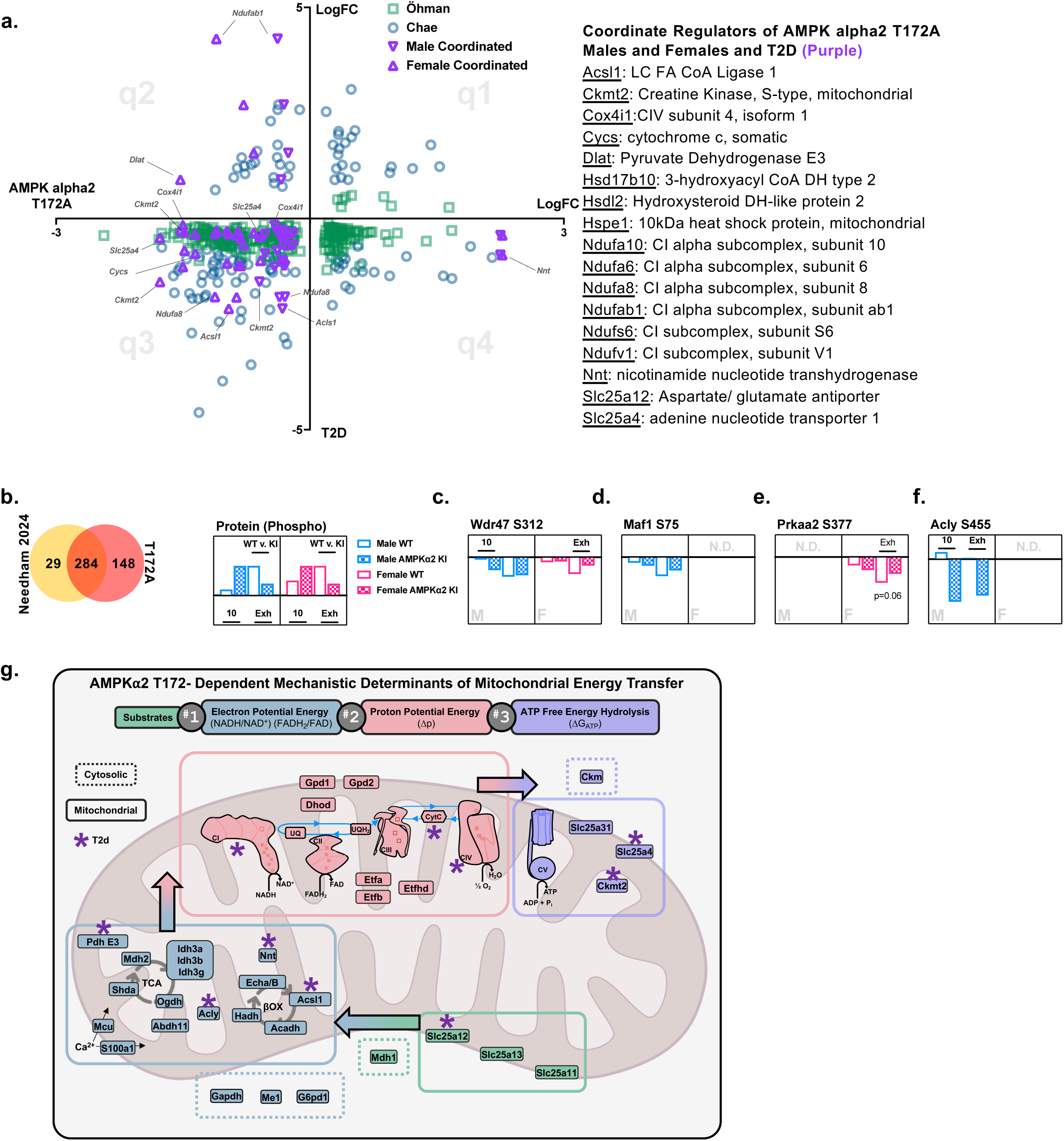
Ampkα2 KI mice share coordinate regulation with type 2 diabetic patient skeletal muscle. (a) Global proteomic results for males and females Ampkα2 mice were compared to two data sets of diabetic human skeletal muscle and plotted in four quadrants (q1-q4) with increased or decreased expression with x-axis representing Ampkα2 T712A KI results and y-axis representing the two human studies (Öhman green □; Chae blue O). Coordinated Ampkα2 KI study results in the Öhman and Chae data sets were presented as male and female data in purple triangles regardless of quadrant agreement and are listed in the table (right) (total 17 proteins with adjusted p<0.05). (b) overlap between Needham 2024^52^ (yellow) and our phosphoproteomic data set (red). (**c-f**) Log2FC of phosphorylation at ATP-citrate synthase (Acly) S455, WD repeat domain 47 (Wdr47) S312, Ampkα2 (Prkaa2) S377, and Maf1 S75 in males and females at 10 min (10) and exhaustion (Exh) (p<0.05). N.D. not detected. (**g**) Mechanisms of mitochondrial energy transfer separated by regulation of substrate (green), electron potential energy (blue), proton potential energy (pink), and free energy of ATP hydrolysis (lilac). All components in the figure were significantly downregulated in global proteomic analysis with adjusted p <0.05. Purple asterisk indicates observed presence in T2D data sets from (**a/b**).

Secondly, phosphoproteomic analysis of insulin-sensitive and insulin-resistant patients with exercise demonstrated significant regulation of 313 unique proteins associated with glucose uptake^52^, overlapping with 284 of the significantly regulated unique genes from our results (**Supplemental Data file 2; Fig 5b**). However, this comparison demonstrated only 4 significantly regulated phosphorylation sites: ATP-citrate synthase (Acly) S455, WD repeat domain 47 (Wdr47) S312, Ampkα2 (Prkaa2) S377, and Maf1 S75 (**Figure 5c-f**). Wdr47 S312, Prkaa2 S377, and Maf1 S75 were significantly altered by exercise at exhaustion, seemingly independent of Ampkα2 T172 regulation. Contrastingly, Acly S455 was dramatically impaired in male KI mice at 10 min of exercise and exhaustion, highlighting the potential importance of this site to Ampkα2 T172 signaling and the development of T2D.

Collectively, these results demonstrate a significant role of Ampkα2 T172 activation in mediating mitochondrial energy transduction, reduction of which putatively leads to metabolic dysregulation in diabetic skeletal muscle, providing strong rationale for targeting Ampkα2 T172 activation to mitigate diabetic pathology (**Figure 5g**; purple asterisk).

## DISCUSSION

Skeletal muscle requires the maintenance of positive energy charge [ATP:ADP] and phosphorylation potential (ΔGp) for its roles in daily function and is highly responsive to physiological changes of bioenergetic demand and proton motive force (Δp) that occur during exercise. It is therefore conceivable that muscle has a sensitive mechanism to detect and react to fluctuation in energetic state, with the purpose of initiating activation of signaling pathways and transcriptional mechanisms that refine bioenergetic form and function. This study reveals that skeletal muscle relies heavily on the energetic bellwether molecule Ampkα2 via its activation by phosphorylation at T172. More than 40 direct Ampk targets have been identified in the literature^36,37^, and the current data set uncovers a litany of novel and vital factors to skeletal muscle metabolism, mitochondrial quantity and quality, and sarcomeric integrity/degradation pathways, amongst others. Predominantly, our results demonstrate that in the absence of Ampkα2 T172 signaling skeletal muscle have diminished metabolic flexibility consequently restricting exercise capacity. Herein, we have mechanistically unveiled key regulatory nodes of Ampk in skeletal muscle at baseline and in response to exercise.

We firstly observed that Ampkα2 KI mice activate glycolysis more rapidly during exercise through several lines of evidence: (1) reduced crossover time and earlier increase of RER during VO2max testing, (2) increased glycolytic flux with exercise through rapid utilization of G1P and accumulation of pyruvate, and (3) greater changes of glycolytic mechanisms observed in phosphoproteomics (e.g., Tbc1d1, Eno3, Pkn2) during exercise. The disconnect we observed between anerobic, and aerobic metabolism appear through pyruvate dehydrogenase (Pdh). Several studies of the role Ampk in exercise indicate a regulatory role of Pdh, with some showing an associated increased expression^53^ and some decreased activity^54^. Our data elucidate that Pdh (Pdha1; Pdhb; Pdhx) mitochondrial pyruvate transport proteins (Mpc1/2) as well as Pdh E2 (Dlat) and E3 (Dld) were all diminished in Ampkα2 KI mice, limiting the incorporation of pyruvate into the TCA cycle.

The flow of energy from electron-rich substrates derived from glycolysis to ATP free energy (ΔGATP) is in the form of mitochondrial energy transduction that is regulated by three nodes^44^: (1) matrix dehydrogenases that generate electron potential energy in the form of NADH and FADH2, (2) the electron transport system maintenance of proton potential energy, and (3) ATP synthesis to maintain ΔGATP. In this regard, impaired metabolic and mitochondrial energy transduction is strongly indicated by the proteomic and metabolomic changes we observed in Ampkα2 KI mice related to each of these three nodes (depicted in **Figure 5g**): (1) Decreased expression of matrix dehydrogenases for cytosolic electron transfer in the form of [NAD^+^]; Hadh), (2) Decreased expression of electron transport chain complexes CI/ CII/ CIII/ CIV for mitochondrial respiration, and (3) Decreased expression of Ckmt2; Ant4; CV directly involved in ATP synthesis and maintenance. Although Ampkα2 KI mice showed dysfunction at each of these steps, our most comprehensive and mechanistic findings point mainly to the importance of nodes 1 and 2. Further studies are required to dissect discrete mechanisms, yet taken together, our results present compelling new evidence for the regulatory role of Ampkα2 T172 activation in governing bioenergetic function and response in skeletal muscle, which dictates exercise capacity.

The primary findings in this study indicates a robust regulatory control of Ampkα2 T172 in mitochondrial quality and function, which has been highlighted in the literature^55^ but infrequently mechanistically ascertained. Our results clearly demonstrate that in the absence of Ampkα2 T172 signaling, mitochondria in skeletal muscle are limited in their capacity to import and incorporate nuclear-encoded proteins into the mitochondria reticulum, construct ETC complexes/supercomplexes, and regulate mitochondrial integrity (**Figure 2d; Figure S4a-d**). These defects may collectively limit metabolic capacity. However, what is unclear from the current analysis is whether the decrements to mitochondrial integrity and electron transport chain assembly proteins are causative to or collateral consequences of diminished aerobic metabolism, as the demand of OXPHOS may enable and expediate formation of machinery that facilitates its function. In this regard, there is some evidence that metabolic demand coordinates the assembly and maturation of individual electron transport chain complexes and larger supercomplexes^56^. Further, Ampk activation through AICAR has recently been utilized to elucidate the mechanisms integrating ETC assembly and efficiency with metabolic cues^57,58^, for which the current data set provides the most comprehensive profiling to date. This specific mechanism may be further informed by a recent study from our group^59^ that demonstrate a novel pool of mitochondria-localized Ampk (mitoAmpk) in skeletal muscle that is responsive to energetic stress.

Our data set has a significant translatable potential as several recent studies have demonstrated that insulin-resistant skeletal muscle has a proteomic signature of decreased OXPHOS machinery, ETC component expression, muscle integrity, and increased proteasomal proteins^60,61^, closely reflected in the expression of these proteins in our Ampkα2 KI mice. Acly S455 phosphorylation was significantly decreased in Ampkα2 KI mice, indicating the importance of this site as downstream of Ampkα2 T172 signaling and vital to metabolism. Further, the 17 key proteins identified have substantial regulatory control over the three nodes of mitochondrial energy transfer: (1) Nnt, Acls1, Slc25a12, Dlat; (2) CI and CIV proteins, cytochrome c; (3) mitochondrial creatine kinase (Ckmt2), Slc25a4 (adenine nucleotide transporter 1), underscoring this mechanism as central to diabetic pathology (**Figure 5**). Although the global proteomic results for Ampkα2 KI highlight significant regulation at nodes 1 and 2, the overlap with T2D muscle points towards the importance of node 3. In this regard, the activation of Ampk with exercise may serve to mitigate diabetic regulation of the mitochondria, as analysis has shown that exercise training activates mitochondrial signaling pathways that directly oppose the effects of diabetes^62^.

In conclusion, our results demonstrate that Ampkα2 T172 activation is critical for both glycolytic and oxidative metabolism and exercise capacity in skeletal muscle. Ampkα2 T172 activation is not only required for the maintenance of metabolic machineries but also for the metabolic flexibility during exercise. Specifically, Ampkα2 activation via T172 phosphorylation dictate the expression of each ETC complex, assembly factors, and components of matrix integrity that are crucial for the maintenance of mitochondrial quality and function. Thus, we have provided foundational experimental evidence with intact protein stoichiometry that Ampkα2 activation is intimately involved in regulating of metabolic energy transfer and oxidative phosphorylation at baseline and during exercise. This is the first study that comprehensively integrates genetic, phenotypic, and multi-omic approaches to provide a high resolution depiction of the multifaceted roles of Ampkα2 T172 activation in skeletal muscle metabolic and contractile functions. Finally, these results position Ampkα2 T172 as a vital regulator that could be augmented in skeletal muscle as a therapeutic targeted against type 2 diabetes and potentially other diseases.

## METHODS

### Knock-In Models & Animal Care

To ascertain the functional role of the Ampk α1 T172 and α2 T172, mutants were developed at the University of Virginia Genetically Engineered Murine Model Core using CRISPR/Cas9 gene editing to replace the Threonine with Alanine at site 172 (T172A) to generate an non-activatable subunit that could not be phosphorylated by upstream kinases. Successful base-pair substitution was confirmed by sequencing and western blot and maintenance of these mouse lines were performed by in-house breeding. Ampk α1 and α2 T172A knock-in mice (C57BL/6 background; 8-12wk/old) and wild-type littermate controls were housed in temperature-controlled (21°C) rooms with 12:12 light-dark cycles and were provided standard chow and water ad-libitum. Male and female mice were used for each outcome described below with number of animals disclosed in each figure legend and are defined by a circle symbol for females and square for males, where appropriate, to facilitate transparency and abide by SAGER guidelines^63^. All experimental procedures were approved by the University of Virginia (#3762) and Fralin Biomedical Research Institute (22-251) Institutional Animal Care and Use Committees.

### Body Composition Measurements

EchoMRI^TM^ (Houston, TX) was utilized for measurements of body composition. Following daily calibration with canola oil system test sample (COSTS) animals were weighed and scanned individually for body composition including a water stage. Outcome measures included fat tissue (g) and lean mass tissue (g). Weights were normalized to bodyweight for fat content or tibia length collected at endpoint for lean mass.

### Metabolic Cage Measurements

Oxymax Comprehensive Lab Animal Monitoring Systems (CLAMS) cages (Columbus Instruments, Columbus, OH) were utilized for indirect calorimetry measures. Following acclimatization an initial set of animals were individually housed for a 24-hour monitoring period followed by 24 hours of monitoring for outcome measures as analyzed by the Oxymax software and included substrate utilization (VCO2, VO2, and respiratory exchange ratio (RER)) and assessment of locomotor activity. Activity is measured by infrared beams at a locomotion axis (X and Y) and rearing axis (Z). Total activity indicates the sum of individual beam breaks during an interval, even at the same coordinates, and can be increased by repetitive beam breaks through activities such as grooming. Ambulatory activity measures the breaking of beams at different coordinates and does not measure repetitive breaking of beams at the same coordinates during an interval.

### Metabolic Tolerance Tests

Metabolic testing included glucose (GTT) and insulin (ITT) tolerance testing with 48 hours in between GTT and ITT. Animals were habituated to handling for three days before beginning the tolerance testing to limit glucose release due to distress. Glucose and insulin were prepared with sterile saline and filter sterilized for 2mg/g and 1U/kg injections respectively. The morning of tests (0900), animals were weighed and individually housed to begin fasting with ad libitum access to water. Following 6 hours (0900- 1500) of fasting, blood glucose was measured at baseline (0hr) utilizing a glucometer (mg/dl; Countour blood glucose meter, Bayer, Leverkusen, Germany) followed by glucose or insulin injection i.p. GTT and ITT performed subsequent blood glucose measures at 30, 60, and 90 minutes and 15, 30, 60 minutes respectively. Following the final measurement, animals were returned to their normal housing cages and provided access to food and water and monitored during recovery.

### Measurements of Exercise Performance and Exhaustive Exercise Procedure

Two separate exercise tests were performed to measure performance via VO2max and Exercise Tolerance (protocols provided in figure 1 and at fbri.vtc.vt.edu/research/labs/yan/protocols.html) **VO2Max** was assessed with a metabolic treadmill (Columbus Instruments) and analyzed via Oxymax software. Animals were acclimated to the treadmill over 3 days immediately prior to testing and included a 10-minute bout at 10m/min 5° incline including shock. Following daily calibration, at the beginning of the light cycle animals were weighed and placed into individual running compartments set to 5° incline. Protocol for VO2Max is demonstrated in figure 1 with a sampling rate of 3 minutes. Animals begin at 0m/min for 2 minutes (stage 1, white bar), followed by 12 minutes of 5m/min “warm up” period (stage 2, gray bar) after which the test begins. Stage 3 consists of 3 minutes of 10m/min followed by 4 minutes of 13 m/min (stage 4, black bar). All subsequent stages (black bar) are 3 minutes and increase by 3m/min with each stage. Exhaustion is defined as an inability or unwillingness to continue running as observed by 5 sequential shocks on the grid at the back of the chamber. When the test is completed the shock grid is turned off and continuous reading of 2 additional cycles is recorded, and the test completed. Animals were returned to their cages and monitored during recovery.

**Exercise Tolerance** or “run to fatigue” protocol was performed on the Columbus Instruments 3/6 treadmill. Treadmill habituation was identical to the VO2Max testing. Stages of the exercise endurance testing are depicted in figure 1. Mice were tested for blood lactate (mmol/L) before beginning (L0) with follow up testing after completing the test (LEnd) to confirm exhaustion. Endurance testing begins with 13.41m/min treadmill speed for 0-30 minutes and increasing to 16.09m/min (30-60min), 18.78m/min (60- 90min), 21.46m/min (90-120min), 24.14m/min (120-150min), and 26.82m/min (>150min) and remained at this speed until exhaustion was confirmed on the shock grid as described above and by LEnd.

**Exhaustive Exercise** was performed identically to the exercise tolerance test with similar determination of endpoint. The first cohort of animals were taken after 10 minutes during which they were immediately sacrificed, and skeletal muscle collected and frozen in metal clamps cooled with liquid nitrogen and then stored at -80°C util time of multi-omic analysis. A second cohort of animals was sacrifice in a similar fashion following determination of exhaustion. Blood lactate was collected as described above to confirm exhaustion.

### Tissue preparation & Western Blot

Mice were euthanized under 2.5% isoflurane at the time of skeletal muscle harvest following an overnight fast. The plantaris, gastrocnemius, soleus, and heart tissues were removed with surgical tools rinsed in 1X PBS, weighed and prepared for different assays accordingly. For western blot, tissues were homogenized in glass homogenizers with protein sample buffer containing 50 mM Tris-HCL, pH 7.4, 0.01% bromophenol blue, 10% glycerol, 1% sodium dodecyl sulfate (SDS), 127 mM 2-mercaptoethanol, and 20 mM dithiothreitol, supplemented with protease inhibitor cocktails and phosphatase inhibitor cocktail tablets (Sigma-Aldrich). Homogenized tissue lysates were then boiled in a heat block at 98 °C for 5 min and stored in -70 °C. Protein concentration was determined by the RCDC protein assay kit (Biorad).

Protein lysates were subjected to sodium dodecyl sulfate-polyacrylamide gel electrophoresis at 100V for 1 hr, and transferred onto nitrocellulose membranes at 80V for 2 hrs. Membranes were probed with the following primary antibodies at a 1:1000 dilution: targeting phospho-Ampkα (Thr172) (CST #2535), Ampk α (CST #2532), phospho-ACC (CST #3661), phospho-ULK1(CST #5869), ULK1 (Sigma-

Aldrich #A7481), GAPDH (CST #2118) and normalized to a muscle common standard (CS) loaded in the middle of the membrane. Secondary antibodies were goat anti-rabbit IR800 and anti-mouse IR680. Membranes were scanned using the odyssey infrared imaging system (LICOR). Proteins were analyzed and normalized to a common protein standard loaded on gel.

### Collection of Mitochondria Enriched Fraction

Plantaris muscle was separated from the soleus and gastrocnemius, weighed, and immediately placed in 1mL of ice-cold mitochondrial isolation buffer (MIB) [BSA (2mg/mL), Sucrose (70mM), Mannitol (210mM), HEPES (5mM), EGTA (1mM), pH 7.1]. At 4°C, muscle was processed with a saw-tooth homogenizer for ∼30 seconds and spun at 4°C for 10 minutes at 800xg to pellet cellular debris. Supernatant was collected and spun again at 9000xg at 4°C for 10 minutes. Supernatant following this spin was collected as cytosolic fraction and the remaining fraction was resuspended in MIB without BSA for quantification of mitochondrial protein by the Bradford method. Protein loading for respirometry was normalized to mg of mitochondrial protein.

### Mitochondrial Respiration

High resolution respirometry was utilized to assess mitochondrial oxygen consumption in an Oroboros O2k (Innsbruck, Austria). We employed the creatine kinase (CK) clamp method during this assay. CK functions to maintain stoichiometric ratio of ATP:ADP during respiration (Creatine + ATP _←_^→^ Phosphocreatine + ADP), which increases the physiological relevance and interpretation of mitochondrial function in contrast to traditional assays that utilize a damaging bolus of ADP. This method provides a discrete and comprehensive measure of mitochondrial oxygen consumption predicated on the regulation of energy demand and energy charge as described previously^44,64^ and depicted in **supplemental figure 4**.

O2k settings remained at 37°C and 500rpm in 0.5mL chambers to daily calibration with buffer D [KMES (105mM), KCl (30mM), EGTA (1mM), KH2PO4 (10mM), MgCl2-6H2O (5mM), 0.05%BSA, pH7.1, solubility factor 0.966] to assess R1 (air saturation, ∼200 µM O2) and closed chamber R0 (zero oxygen) with sodium hydrosulfide (Sigma S1256). Chambers were then washed with DI water (5 minutes for 5 cycles) before beginning experiments. Oxygen flux (*J*O2; pmol/sec) was assessed in buffer D supplemented with creatine monohydrate (5mM) and 20mg of mitochondrial protein. State 2 respiration (leak state) was assessed with pyruvate (5mM), malate (2.5mM), and octanoyl carnitine (0.2mM) followed by state 3 (maximal respiration; ΔGATP -12.94 kCal/mol) with creatine kinase (20mM), phosphocreatine (1mM), and ATP (5mM). Cytochrome c (0.005mM) was used to assess mitochondrial integrity with a threshold of 15% increase in respiration. Sequential PCr titrations were then added at 1mM (ΔGATP -13.38 kCal/mol), 2mM (ΔGATP -13.71 kCal/mol), 4mM (ΔGATP -14.12 kCal/mol), and 6mM (ΔGATP -14.45 kCal/mol).

Calculations of ΔGATP were performed using an online calculator (provided here^44^) under the conditions of 37°C, 170mM ionic strength, 5mM creatine, 10mM phosphate, and pH 7.1. The plotting of *J*O2 vs. ΔGATP across the linear force-flow relationship allows for calculation of mitochondria “conductance”. Conductance measures the inverse of resistance (∼1/R) of the mitochondrial proton motive force (Δp; voltage) in relationship to Ohm’s law (I=V/R) where current (I) measures *J*H+.

### LC-MS/MS Proteomics & Phosphoproteomics

Muscle tissue samples from male and female WT and KI mice were collected before exercise (i.e., sedentary control, SED), after 10-min running (10 min) and after exhaustion (Exh.) and stored at - 80°C until undergoing processing. Sample processing for global proteomics and phosphoproteomics analysis was conducted as previously described^65^ with a several modifications. Briefly, frozen muscle tissue samples were minced into small pieces on a pre-chilled aluminum tray with dry ice. ∼ 25 mg of tissue were collected and homogenized in cold lysis buffer (50 mM Tris, 8 M urea, 75 mM NaCl, 1 mM EDTA, 2 µg/mL aprotinin, 10 µg/mL leupeptin, 1 mM PMSF, 10 mM NaF, 1% phosphatase inhibitor cocktail 2, 1% phosphatase inhibitor cocktail 3, pH 8.0) on a pre-chilled bead beater using 2-min cycle. Samples were then centrifuged at 13,000xg at 4 °C for 10 min to remove debris. Extracted protein was quantified by BCA assay. The same quantity of protein was aliquoted and reduced with 5 mM DTT for 1 Ampk shaking on a thermomixer at 37 °C, 1000 rpm. Proteins were then alkylated with 20 mM iodoacetamide with shaking in dark at room temperature, 1000 rpm for 45 min. Samples were diluted 4 fold, then digested with trypsin (enzyme to protein ratio = 1: 50) for 1hr, shaking at 1000 rpm, 37°C. A fresh aliquot of trypsin (enzyme to protein ratio = 1: 50) was added after the samples were diluted 2 folds. Then the digestion was conducted at 37°C 1000 rpm overnight. The digest was desalted by Sep-Pak C18 SPE cartridges (Waters, Milford, MA). Clean peptides were quantified by BCA assay. 250 µg of peptides from each sample was aliquoted and concentrated in a speedvac to completely dry to be used for TMT labeling.

Peptides were resuspended in 500 mM HEPES (pH 8.5) at 5 μg/μL. Three replicates of male or female muscle tissue samples from each condition (WT-SED, WT-10 min, WT-Exh., KI-SED, KI-10 min, KI-Exh.) were labeled with two TMT18 plexes. TMT reagents were resuspended in anhydrous acetonitrile at 20 μg/μL and added to each sample at a 1:2.5 (peptide:TMT) ratio. Labeling was conducted at 25 °C, 850 rpm for 1hr shaking on a thermomixer. Then the reaction was quenched by hydroxylamine. Samples from each plex were combined and concentrated in a speedvac, followed by C18 SPE cleanup. The clean TMT-labeled samples were then fractionated into 12 fractions using high pH reversed phase separation. 5% of each fraction was used for global proteomics analysis, the remained fractions were subjected to immobilized metal affinity chromatography (IMAC) phosphoenrichment using freshly prepared Fe^3+^-NTA-agarose beads.

Both global and phosphopeptide fractions were analyzed using Waters nanoAcquity UHPLC system (with 20 cm x 75 um i.d. 1.9-um column packed in-house with Waters BEH C18 silica) coupled to a Orbitrap Fusion Lumos (Thermo Scientific, San Jose, CA) with a 120-min LC gradient. Positive ion mode spray voltage was set at 2.2 kV. Full MS spectra were recorded at resolution of 60K with scan range 350 – 1800 m/z. Automated gain control (AGC) value was set 4e5. MS/MS was acquired in data dependent mode (DDA) at a resolution of 50K, AGC of 1e5. Isolation window was 0.7 m/z. High-energy collision dissociation (HCD) with a normalized collision energy setting of 30% was used. Dynamic exclusion time was set at 45s.

### Untargeted Metabolomics

Muscle tissue samples from male and female wild-type (WT) and knock-in (KI) mice were collected as described above. For extraction, tissues were thawed on ice, and approximately 750 µL of a cold chloroform: methanol (1:2) mixture was added, along with four steel balls (Fisher Brand; diameter 2.4 mm). Tubes were plunged into liquid nitrogen for 5 minutes and vigorously shaken in a Fisher Brand Bead Mill 24. Tubes were then vortexed and shaken at 900 rpm for 30 minutes at 4°C in a temperature-controlled thermal shaker. After adding 400 µL of water, samples were vortexed, and the upper aqueous phase was recovered as the metabolite mixture. Two hundred microliters of the extract were dried overnight in a SpeedVac and reconstituted in 200 µL of 0.1% formic acid (FA) containing the 100X Metabolomics QReSS™ Kit (Cambridge Isotopes, MSK-QRESS-KIT). A 10 µL aliquot from each tube was removed to create a pooled quality control (QC) sample, which was injected at the beginning and end of the mass spectrometry (MS) sequence. Additional QC samples were injected after every five sample injections.

LC-MS data acquisition was performed using a fully automated AcquireX Intelligent Data Acquisition Workflow with additional instrument parameters, data acquisition and data processing workflow is based on a previous publication^66^. We first generated an exclusion list was generated from a reagent blank sample to determine the background. A pooled quality control (QC) sample was injected for feature detection and component assembly to create an inclusion list. Using the inclusion list, a series of iterative data-dependent acquisition (DDA) injections were performed, with each injection informed by the previous one. Precursors from the inclusion list were fragmented, and once detected, they were automatically transferred to the exclusion list. This approach minimized redundant fragmentation and maximized relevant spectra and metabolite annotation. Samples were analyzed on a Thermo Orbitrap IDX Tribrid MS coupled to a Thermo Vanquish UHPLC. Metabolite separation was achieved using a Waters BEH C18 column (Waters Corp.; 2.1 × 150 mm, 1.7 µm) maintained at 30°C. Samples were analyzed in both positive and negative ionization mode.

### Differential Analysis

Global proteomics and phosphoproteomics were searched against protein sequences from the 2023 UniProt database with the MS-GF+ tool^66,67^. After removal of contaminant proteins, correction for isotope selection error, and phosphosite localization with the Ascore method^68^, peptide-spectrum matches (PSMs) were filtered to limit the peptide-level FDR to 1%. This was followed by a 1% protein-level FDR filter for global proteomics, or the mapping of phosphopeptides to phosphosites and limiting the phosphosite-level FDR to 1%. Decoy PSMs were removed and the parsimonious set of proteins was determined by preferentially assigning peptides to the protein with the highest number of total peptide identifications. For phosphoproteomics, phosphosites were first assigned to any proteins detected in the global proteomics results before assigning them to the protein with the most peptide identifications. The MASIC intensity data was then combined with the processed MS-GF+ search results and intensities were aggregated at the level of proteins or phosphosites and log2 transformed^69^. Since males and females were measured in different TMT plexes, they were separated, and any proteins or phosphosites that were not measured in all samples for a particular sex were removed. Then, samples were normalized by subtracting their median intensities to remove differences due to sample loading and each protein or phosphosite was median-centered.

For the global proteomics, phosphoproteomics, and metabolomics datasets, we fit a linear model that included a single predictor with all combinations of group (KI and WT) and timepoint (SED, 10 min, EXH) as categories using the lmFit function from the limma R/Bioconductor package^70–72^. For global proteomics, the presence of several high variance samples led to the use of sample-specific quality weights; these weights, generated automatically with the arrayWeights function from limma, reduce the contribution of samples with high variance without reducing the available degrees of freedom^74^. Contrasts were constructed to test for differences between the SED KI and SED WT groups in global proteomics. The phosphoproteomics contrasts compared the 10 min and EXH timepoints to the SED timepoint within both KI and WT groups, in addition to testing for differences in the EXH vs. SED response between the KI and WT groups: (KI.EXH - KI.SED) - (WT.EXH - WT.SED). Lastly, the metabolomics contrasts included the four trained vs. SED contrasts from the analysis of the phosphoproteomics data and the SED KI vs. SED WT comparison that was tested for the global proteomics data. For all omes, the eBayes function from limma was used to squeeze the residual variances toward a global trend that was made robust to the presence of any outlying features^71,73^. P-values were adjusted across the trained vs. SED contrasts and separately for the other contrasts using the method of Benjamini and Hochberg to control the false discovery rate (FDR)^74^. P-values were adjusted across the trained vs. SED contrasts and separately for the other contrasts using the method of Benjamini and Hochberg to control the false discovery rate (FDR)^76^.

### Enrichment Analysis

The global proteomics, phosphoproteomics, and metabolomics differential analysis results were each transformed into matrices of z-scores (calculated from moderated t-statistics) separated by sex with gene symbols (proteomics), singly-phosphorylated sites (phosphoproteomics), or Reference Set of Metabolite Names (RefMet) metabolite identifiers as rows and contrasts as columns^75^. In the phosphoproteomics matrices, only the exercise vs. SED contrasts were utilized. To resolve situations where multiple proteins map to the same gene symbol or separation of multiply-phosphorylated phosphosites results in multiple rows for the same site, the feature with the most extreme z-score (smallest p-value) was selected per contrast. If any proteins did not map to a gene symbol, the protein identifier was used to avoid the unnecessary removal of data. This procedure resulted in a total of 3,365 genes, 17,636 phosphosites, and 452 RefMet metabolite identifiers from either sex.

Version 2023.2 of the mouse Molecular Signatures Database (MSigDB), particularly the C5 collection of 10,662 Gene Ontology terms, was selected for the analysis of the global proteomics data^76–79^. Of these terms, 3,042 contained at least 10 genes that overlapped with the set of 3,365 genes measured in the proteomics data. These ∼3000 gene sets were then clustered according to their Jaccard similarity coefficients using the clusterSets function from the TMSig R package to identify groups of highly similar gene sets^80^. The largest gene set in each cluster was selected as the representative for testing. If two or more gene sets had the same maximum size, the set with the highest proportion of original genes was selected; further ties were broken by selecting the set with the shortest description (often, the most general term) and then the first set alphabetically. This removed 194 redundant gene sets.

The “Kinase_Substrate_Dataset” file was downloaded from PhosphoSitePlus (v6.7.1.1), and the data was filtered to mouse kinases and substrates^81^. The phosphosites, formed by concatenating the SUB_GENE and SUB_MOD_RSD columns, were then grouped by their known kinases. Only 26 out of 239 possible kinase sets contained at least 3 phosphosites that overlapped with the set of 17,636 phosphosites from the male or female phosphoproteomics datasets.

Metabolite subclasses from the RefMet database were selected for the analysis of the metabolomics data^75^. Only 9 subclasses contained at least 10 metabolites that overlapped with the set of 452 measured metabolites.

Gene sets, kinase sets, and RefMet subclasses were analyzed with the parametric version of the pre-ranked correlation adjusted mean rank (CAMERA-PR) test^43^. This test is a modification of the two-sample t-test that accounts for correlation between molecules within sets to better control the type I error rate. The analysis was carried out on the z-score matrices with the cameraPR.matrix function from TMSig^80^. As with the differential analysis results, p-values were adjusted across contrasts using the BH method^74^.

### Statistical Analysis (Non-Omic Results)

Data are represented as mean ± SEM. In analyses where one variable was present data were analyzed with a student’s t-test. Two-way ANOVA testing was performed for several outcomes where multiple variables were present to demonstrate main effects. Statistical significance was established *a priori* as p < 0.05 or p <0.01 where indicated. Data presented as mean ± SEM. Statistical analysis performed by t-test between groups: p < 0.05 (*),p < 0.01 (**),p < 0.001 (***), and p < 0.0001 (****).

## Supplemental figure captions

**Figure S1:**
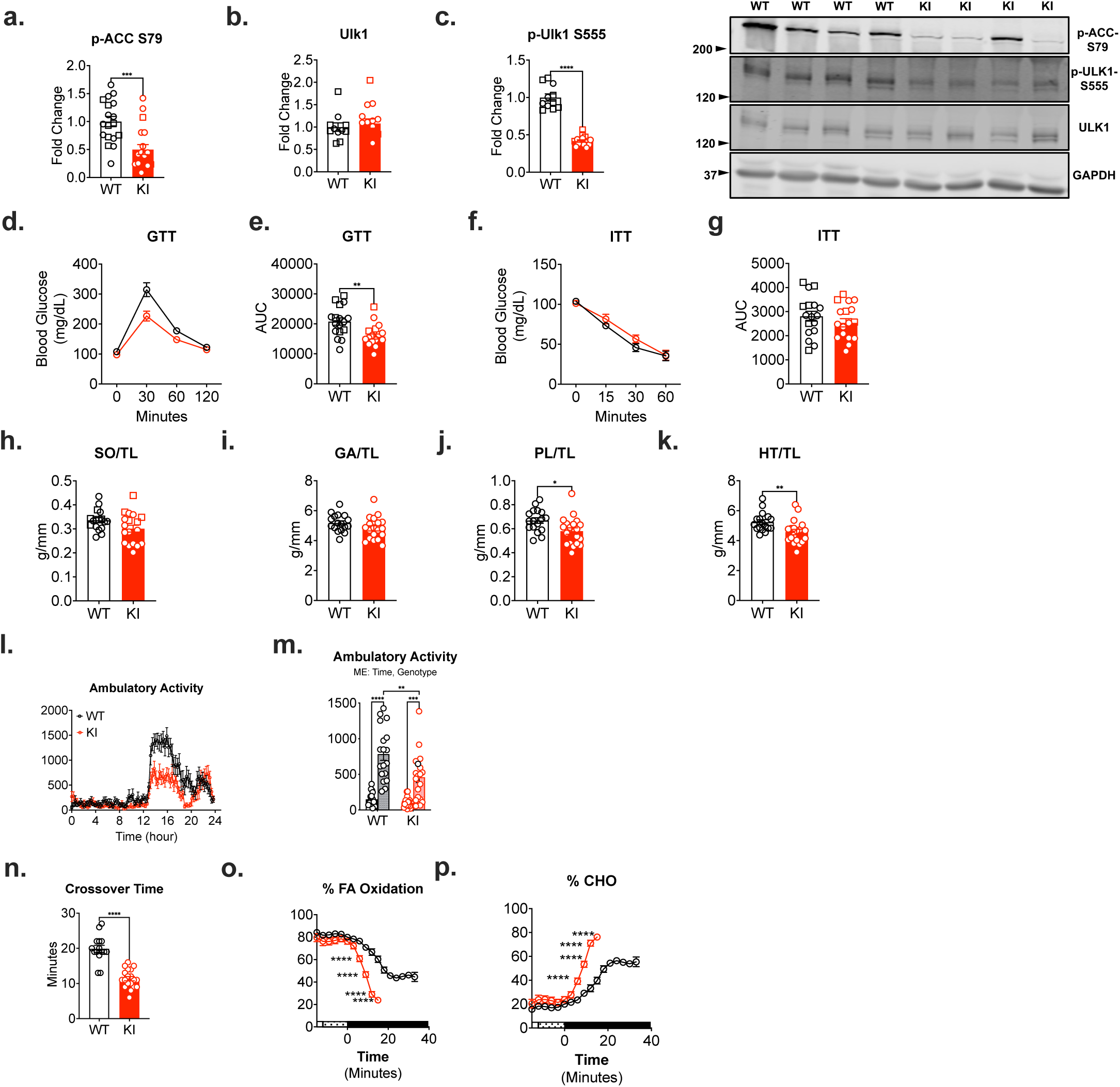
Supplement: Ampkα2 KI mice demonstrate altered metabolic regulation and limited exercise capacity. (**a-c**) Western blot confirmation of classical Ampk signaling targets pACC (phosphorylated acetyl-CoA carboxylase) and total and pS555-Ulk1 and representative image. (**d-g**) Glucose (GTT) and insulin tolerance testing (ITT) and area under the curve (AUC) analysis. (**h-k**) Skeletal muscle weights (g) at the time of sacrifice: soleus (Sol), gastrocnemius (GA), plantaris (PL), and heart (HT) normalized to tibia length (TL; mm). (**l/m**) Metabolic cage measurements (Columbus Instruments) were taken over 24 hours for ambulatory activity. Comparisons made between the light/ resting (0700-1900; time 0-12 hours) and dark/ active (1900-0700; time 12-24 hours) cycles. (**n-p**) VO2Max testing evaluation of crossover time to reach anaerobic threshold as demonstrated by percent fatty acid (%FA) oxidation and percent carbohydrate (%CHO) oxidation. Males (n=6 WT; n=7 KI) represented in squares and females (n=11 WT; n=12 KI) in circles for all outcomes; circles represent genotype average for **d**/**f**/**l**/**o**/**p**. Data presented as mean ± SEM. Statistical analysis performed by t-test between groups. Two-way ANOVA performed for **m** for main effects (ME) of time and genotype. Significance indicated as p < 0.05 (*), p < 0.01 (**), p < 0.001 (***), and p < 0.0001 (****), ns= not significant.

**Figure S2:**
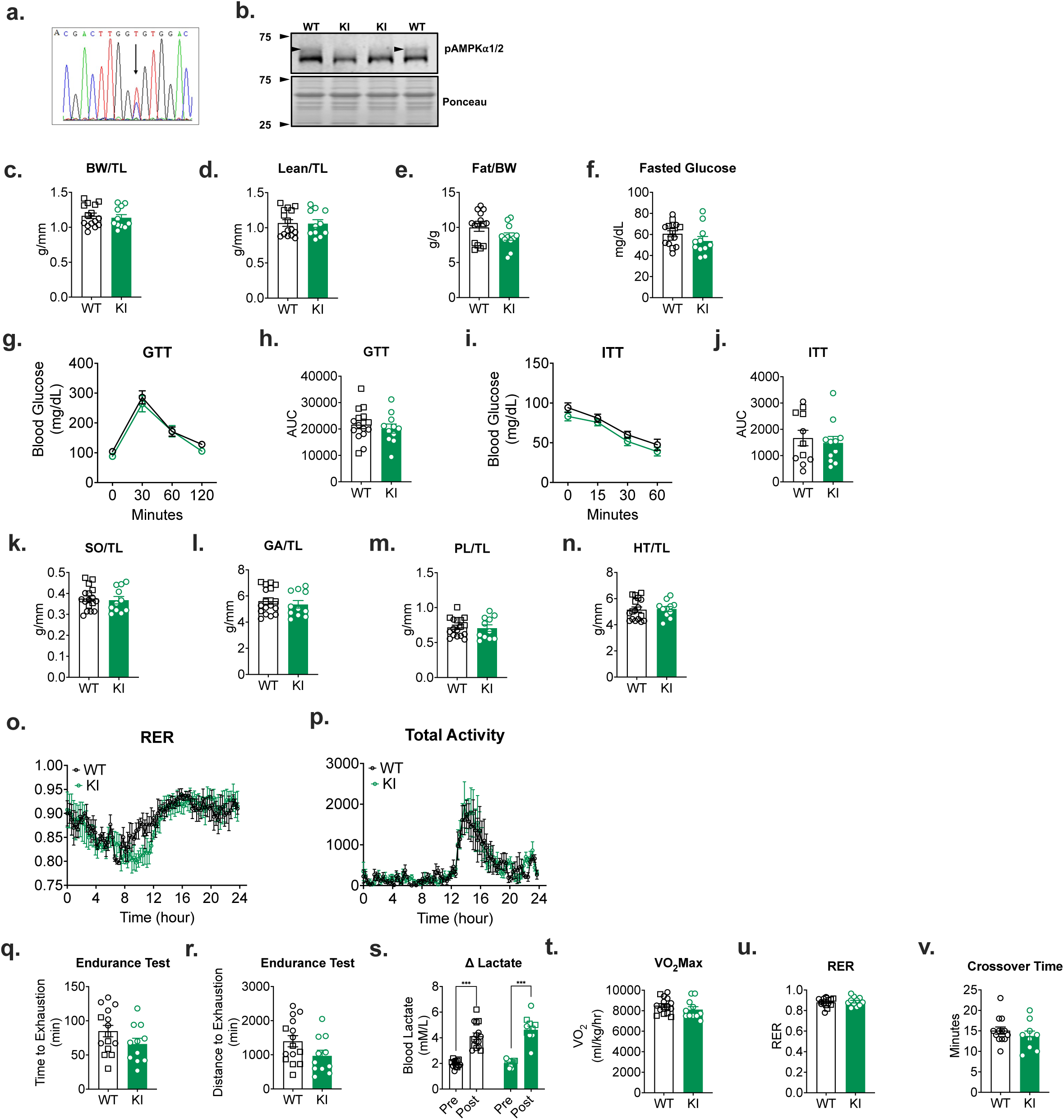
Supplement: Ampkα1 KI mice present normal metabolic and exercise capacity. (**a/b**) Genotype was confirmed with sequencing and western blot for the T>A mutation at Ampkα1 T172 site. (**c-e**) Body composition analysis (EchoMRI) evaluated bodyweight (g) as well as lean mass (g) normalized to tibia length (TL; mm) and fat mass (g) normalized to bodyweight (g). (**f**) Fasted glucose measurements were taken after an overnight fast (1900-0700). (**g-j**) Classical glucose (GTT) and insulin tolerance testing (ITT) and area under the curve (AUC) analysis. (**k-n**) Skeletal muscle weights (g) at the time of sacrifice: soleus (Sol), gastrocnemius (GA), plantaris (PL), and heart (Ht) normalized to tibia length (TL; mm). (**o/p**) Metabolic cage measurements (Columbus Instruments) were taken over 24 hours for total activity and respiratory exchange ratio (RER). Endurance testing and VO2Max testing performed to evaluate exercise capacity (protocol indicated in figure 1). (**q-s**) Endurance testing evaluated in minute and meters and confirmed exhaustion by pre-post lactate measurements. VO2Max testing evaluation of VO2Max, RER, and crossover time to reach anaerobic threshold. Males (n=4-7 WT; n=4-6 KI) represented in squares and females (n=8 WT; n=5 KI) in circles for all outcomes; circles represent genotype average for **g**/**i**/**o**/**p**. Black/ white indicates WT and green indicates Ampkα1 KI. Data presented as mean ± SEM. Statistical analysis performed by t-test between groups. Significance indicated as p < 0.05 (*), p < 0.01 (**), p < 0.001 (***), and p < 0.0001 (****), ns= not significant.

**Figure S3:**
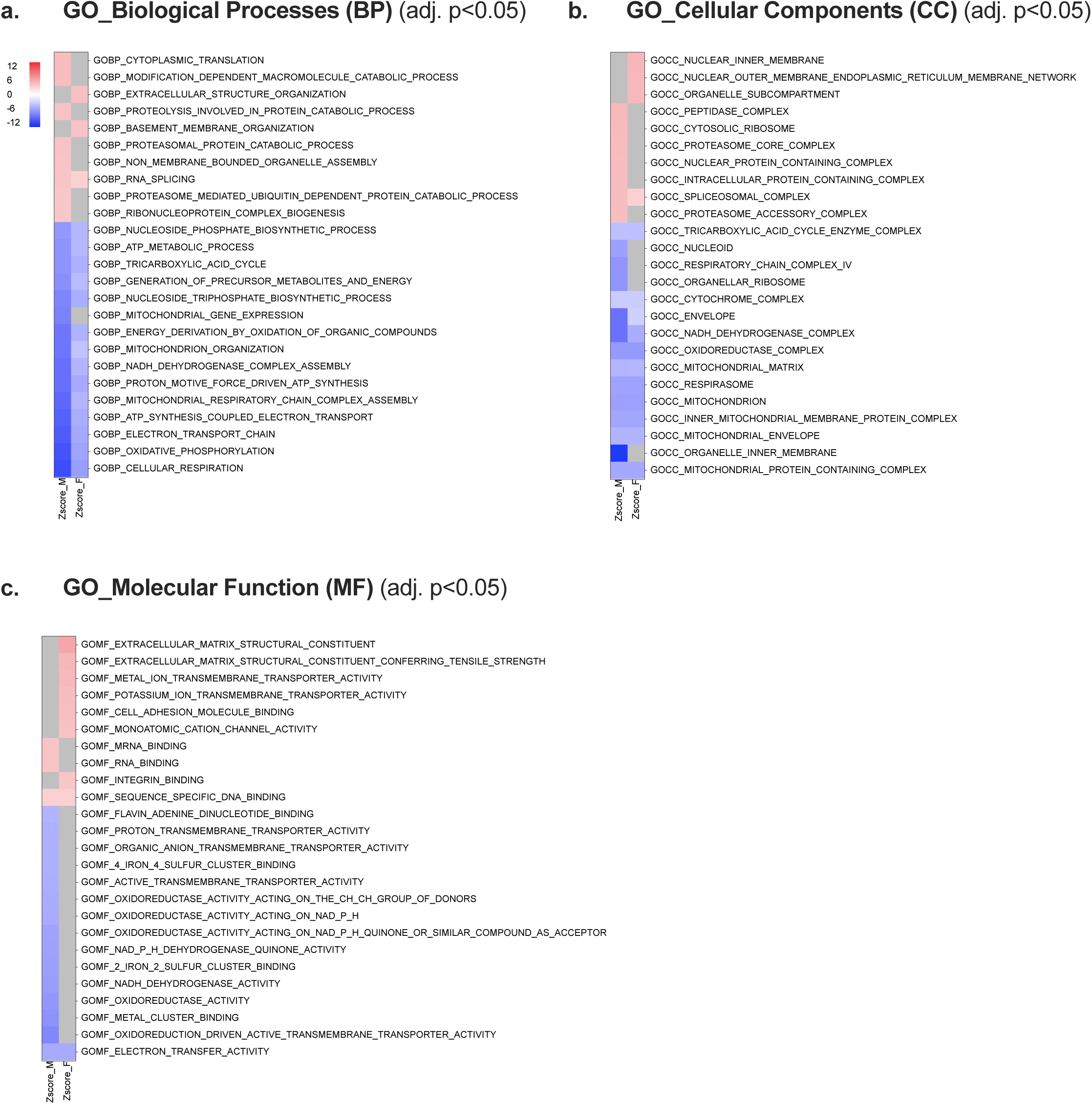
Supplement: Results of global proteomics separated by gene ontology (GO) terms. (**a-c**) Top 10 significantly upregulated (red) and top 15 significantly downregulated (blue) terms for GO terms (GO biological processes (BP); cellular components (CC) and molecular function (MF) represented by Z score. Gray box indicates no result. All results are adjusted p<0.05. Full results present in Supplemental data file 1.

**Figure S4:**
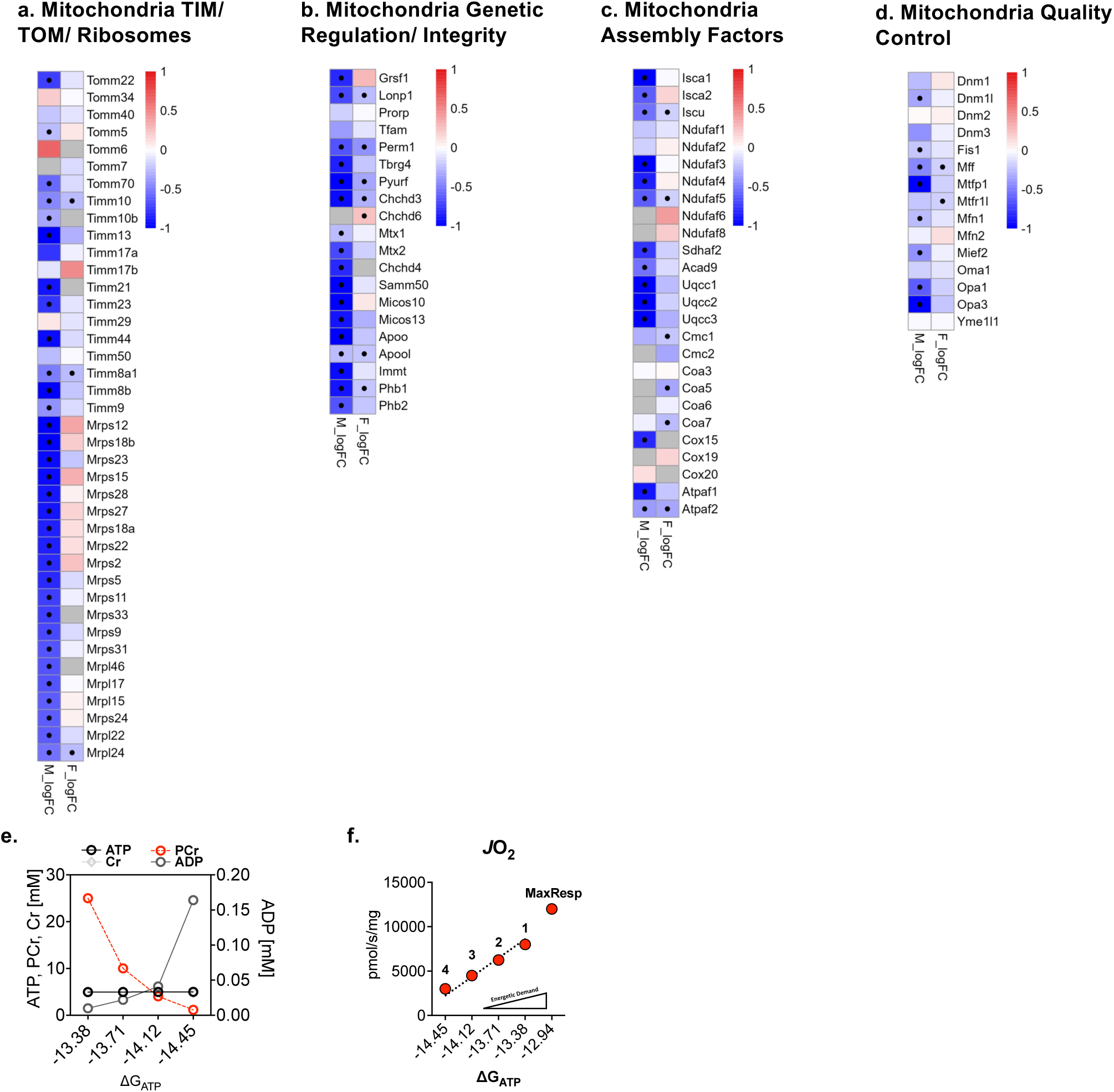
Supplement: Enrichment analysis of global proteomics with MitoCarta overlay, creatine kinase clamp method illustration. Full results present in S**upplemental data file 1**. Red indicates increased expression and blue decreased with • representative of adj. p < 0.05 difference within heatmap by t-test (WT n=6; KI n=6). (**a**) Global proteomic processes related to mitochondrial translocase of inner membrane (TIM), translocase of outer membrane (TOM), and representative mitochondrial ribosome proteins (Mrps/Mrpl). (**b**) regulation of genetic processes within the mitochondria and mitochondrial gene transport/processing. (**c**) mitochondria/ electron transport chain assembly factors. (**d**) mitochondrial quality control. (**e/f**) Representative concentrations of ATP, phosphocreatine (PCr), creatine (Cr), and ADP during the creatine kinase clamp evaluation of respiration and titration of PCr to represent conductance over a range of energy demands. Statistical analysis performed by t-test between groups: adjusted p < 0.05 (•).

**Figure S5:**
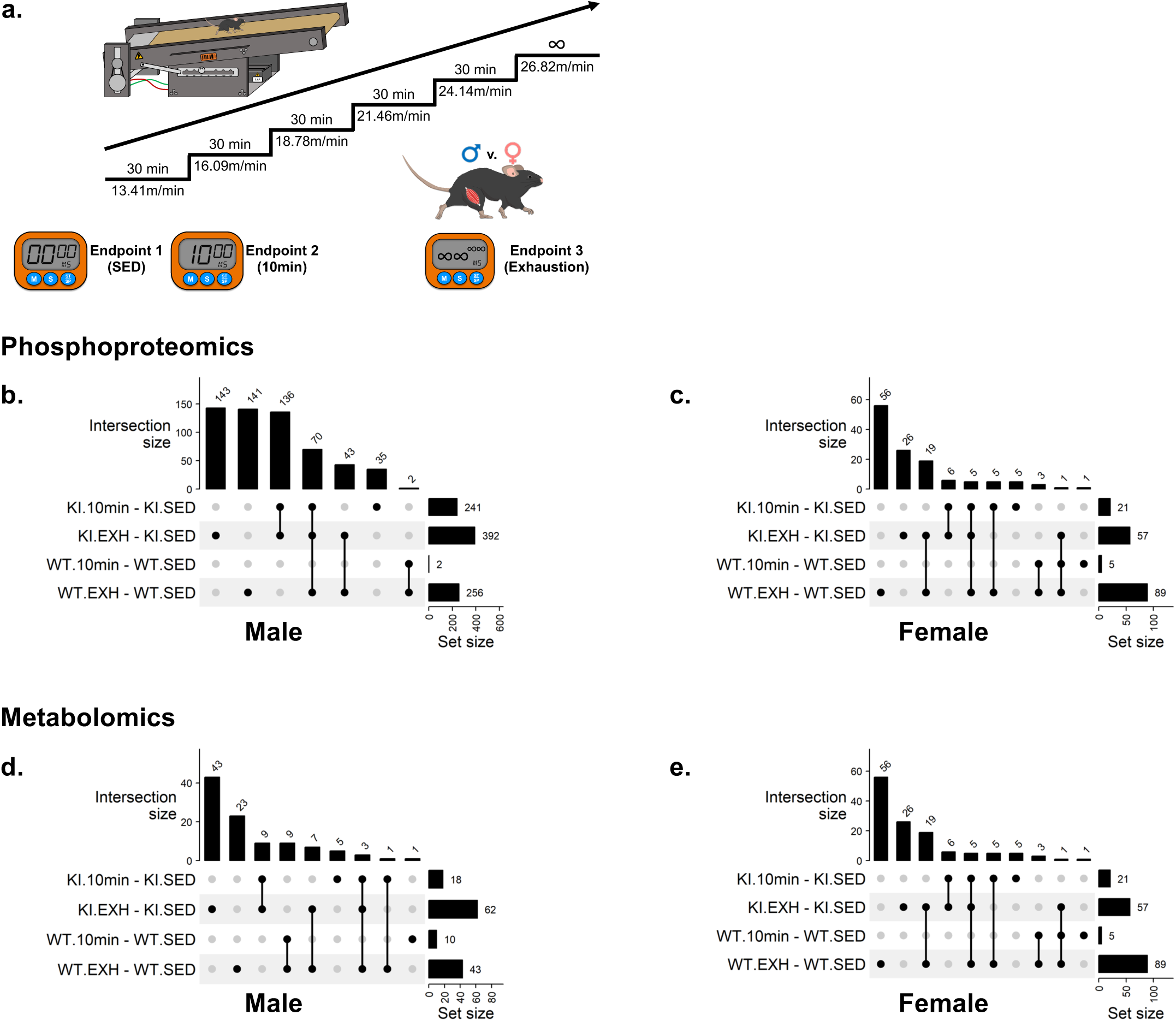
Supplement: Ampkα2 T172 dictates the phosphoproteomic and metabolomics response to exercise within skeletal muscle. (**a**) representative method of the exhaustive exercise protocol with endpoints at 0 minutes of exercise (SED), 10 minutes of exercise (10) and exhaustion (∞) (WT n=6; KI n=6). (**b/c**) Upset plots of statistically significant male and female phosphoproteome following 10 minutes of exercise and at exhaustion separated by genotype. (**d/e**) Upset plots of statistically significant male and female metabolome following 10 minutes of exercise and at exhaustion separated by genotype.

**Figure S6:**
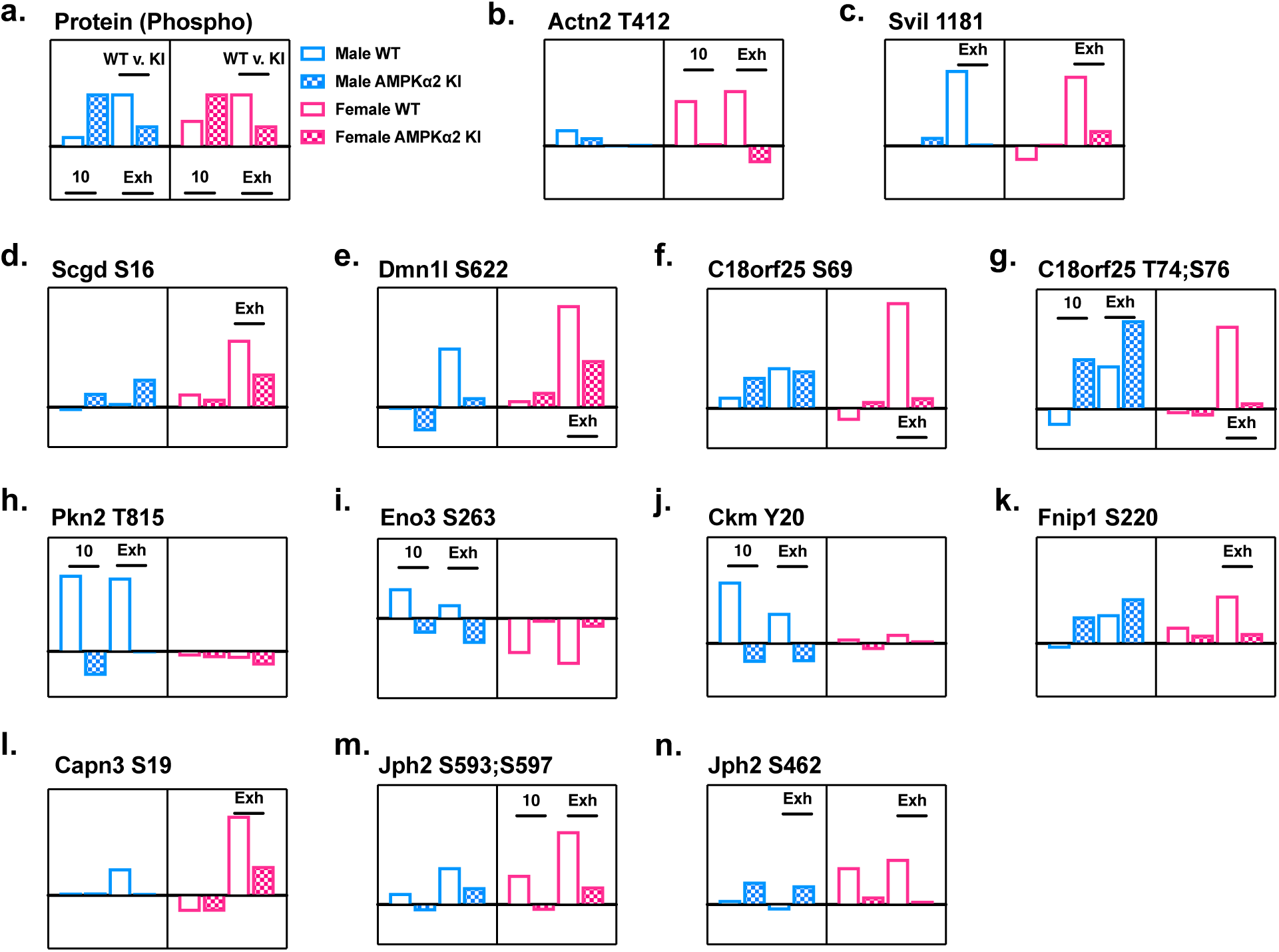
Select phosphoproteomic annotations from Ampkα2 T172 phosphoproteomic response to exercise within skeletal muscle. (**a**) WT (open bar) and KI (checkered bar) mice were divided into three treatments (1) sedentary, (2) exercise for 10 minutes (10min) and (3) exercise to exhaustion (Exh) groups further by sex for male (M; blue) and female (F; pink). Bars above timepoints indicate significant differences between WT and Ampkα2 KI mice at 10 minutes or exhaustion. Complete data provided in **supplemental data file 2**. (**b- n**) Log2FC of phosphorylation sites representing significantly increased phosphorylation in males and/or female WT (adj. p<0.05) *and* a significant decrease compared to KI (p<0.05) at exhaustion. Actn2 actinin2; Svil supervilin; Scgd sarcoglycan delta; Dnml1 dynamin like protein; Pkn2 protein kinase N2; Eno3 enolase 3; Ckm creatine kinase muscle; Fnip1 follicular interacting protein 1; Capn3 calpain 3; Jph2 junctophilin 2.

**Figure S7:**
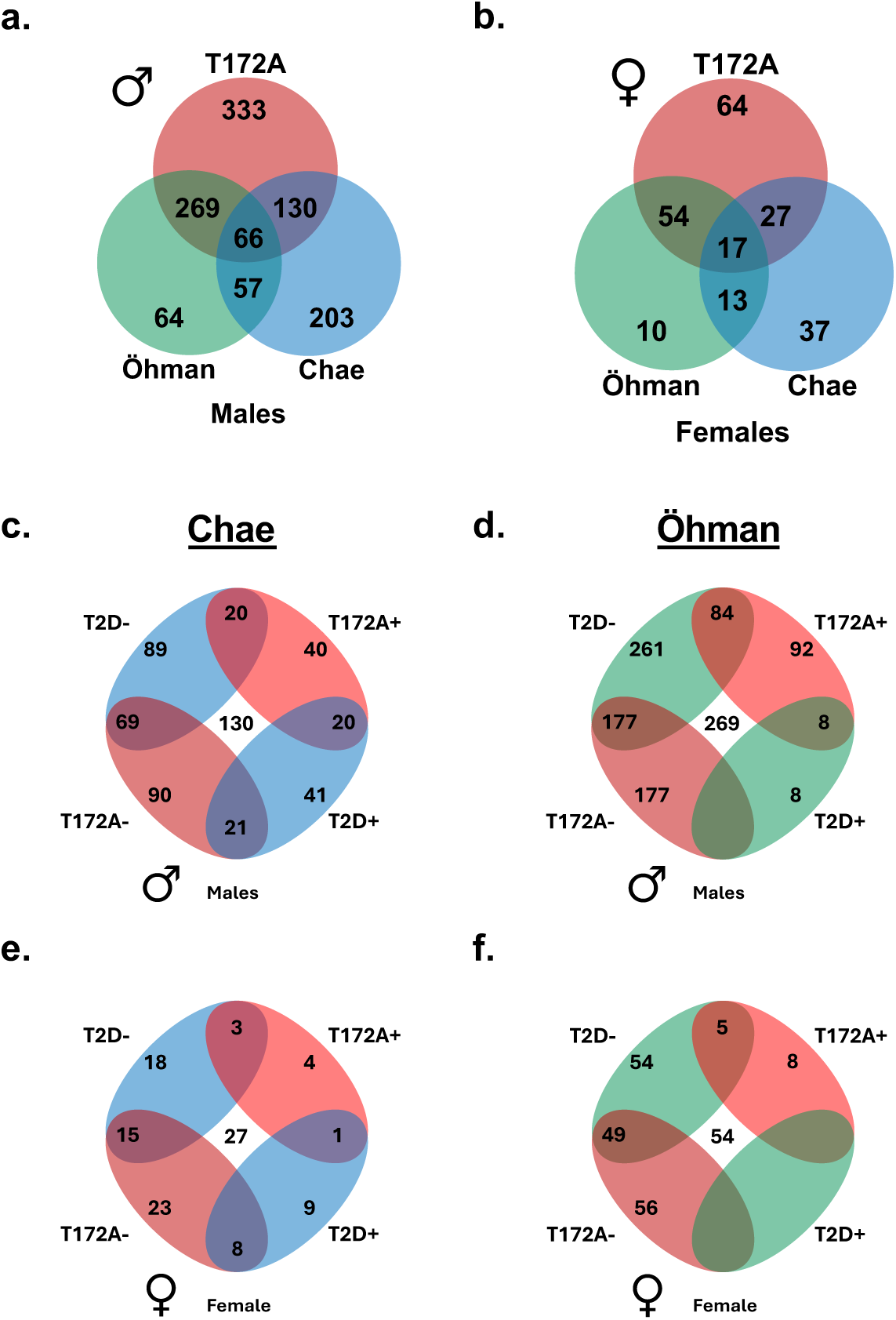
Supplement: Global proteomics of Ampkα2 T172A KI shares coordinate regulation with type 2 diabetic patient skeletal muscle. (**a/b**) Males (M) and females (F) from our data set (red) share 66 and 17 individual proteins respectively with both Öhman (green) and Chae (blue) the diabetic data sets. The blue and green circles that do not overlap represent the number of genes that were present in the respective studies but significantly altered in the Ampkα2 T172A data set (Chae 203M; 37F/ Öhman 62M, 10F). (**c-f**) Where overlap was observed (Chae 130M; 27F/Öhman 269M, 54F) results were distributed in the 4-way matrices dependent on agreement between positive (+) and negative (-) regulation of T172A genes and the diabetic data sets.

## Supplemental File 1: Global Proteomics

- File name: S1_Ampk_global_DA_CAMERA_male_and_female.xlsx

## Supplemental File 2: Phosphoproteomics

- File name: S2_Ampk_phospho_DA_male_and_female_Ampk_delta10min_deltaExh_XL.xlsx

## Supplemental File 3: IPA analysis

- File name: S3_Ampk_phosphoprot_IPA_results.xlsx

## Supplemental File 4: Metabolomics

- File Name: S4_Ampk_metab_DA_male_and_female.xlsx

## Resource Availability

The data generated in this study are available from the corresponding author upon request, have been deposited in publicly available and are available in supplemental data files provided here. Databases can be referenced under the manuscript title at open science framework (www.OSF.io)), MassIVE (massive.ucsd.edu), and ProteomeXChange. Request for further information and resources should be directed to and will be fulfilled by the corresponding author.

## Author Contributions

**(CRediT** https://credit.niso.org/): All authors have approved of the submitted version of the manuscript.

Conceptualization: RNM, WJQ, ZY

Data curation: RNM, TS, XL, JY, ARB, MAG, MGJ, WS, NW, WJQ, ZY

Formal analysis: RNM, TS, XL, JY, NW, ARB, HB, WJQ, ZY

Funding acquisition: RNM, ZY

Investigation: RNM, GM, TS, XL, JY, WS, ARB, MAG, MGJ, NW, HB, YG, XM, MZ, WJQ, ZY

Methodology: RNM, GM, TS, XL, NW, WJQ, ZY

Project administration: RNM, WJQ, ZY

Resources: WJQ, ZY

Software: TS, XL, NW, MAG, MGJ, HB, WJQ, ZY

Supervision: WJQ, ZY

Validation: RNM, GM, TS, XL, ARB, MAG, MGJ, NW, MZ, WJQ, ZY

Visualization: RNM, TS, XL, NW, ARB, HB, WJQ, ZY

Writing – original draft: RNM, GM, ZY

Writing – review & editing: RNM, GM, TS, XL, JY, WS, MAG, ARB, MGJ, NW, HB, YG, XM, MZ, WJQ, ZY

## Declaration of generative AI in scientific writing

No components of this manuscript writing, analysis, interpretation or representation were prepared with the use of generative artificial intelligence.

